# Isotopic tracing of [^13^C_6_]*scyllo*-inositol uncovers its incorporation into phosphatidylinositols in mammalian cells

**DOI:** 10.64898/2026.04.07.716873

**Authors:** Meike Marie Amma, Laxmikanth Kollipara, Peter Schmieder, Adolfo Saiardi, Sven Heiles, Dorothea Fiedler

**Affiliations:** Leibniz-Forschungsinstitut für Molekulare Pharmakologie (FMP), Robert-Rössle-Str. 10, 13125 Berlin, Germany; Institut für Chemie, Humboldt-Universität zu Berlin, Germany, Brook-Taylor-Str. 2, 12489 Berlin, Germany; Leibniz-Institut für Analytische Wissenschaften-ISAS e.V., Dortmund, Germany; Faculty of Chemistry, University of Duisburg-Essen, Essen, Germany; Laboratory for Molecular Cell Biology, University College London, London WC1E 6BTU.K

**Keywords:** [^13^C_6_]*scyllo*-inositol, isotopic tracer, phosphatidylinositol, metabolomics, lipidomics, mass spectrometry, NMR, HILIC

## Abstract

Inositols are a family of cyclic sugar alcohols comprising nine stereoisomers. *Myo*-inositol is the most abundant isomer found in humans and has been studied most extensively. It plays an important role in osmoregulation and is incorporated into membrane-anchored phosphatidylinositols. *Scyllo*-inositol is the second most abundant inositol isomer in the human brain and aberrant concentrations are associated with various diseases; however, its biological functions remain poorly understood. Here, the development and application of [^13^C_6_]*scyllo*-inositol as an isotopic tracer to study its metabolism is reported. A concise and robust synthetic route was established to obtain [^13^C_6_]*scyllo*-inositol from [^13^C_6_]*myo*-inositol in good yield. The uptake of [^13^C_6_]*scyllo*-inositol and responses of endogenous inositol isomers were measured in multiple cell lines by HILIC-MS/MS, showcasing the advantages of isotopic tracing. [^13^C_6_]*scyllo*-inositol proved to be a versatile isotopic tracer, when coupled with MS-based lipidomics and 2D NMR experiments. These experiments provide evidence that *scyllo*-inositol is incorporated into phosphatidylinositols in different cell lines. The results suggest a previously underappreciated role of *scyllo*-inositol in mammalian cells. The utilization of [^13^C_6_]*scyllo*-inositol will help to elucidate the role of *scyllo*-inositol metabolism in healthy and diseased states.

**Significance:** *Scyllo*-inositol is a cyclic sugar alcohol found predominantly in the human brain. Changes in its concentration are associated with different diseases, and *scyllo*-inositol has been investigated as a potential drug against Alzheimer’s disease in clinical trials. However, its metabolic fate in mammalian cells is not well understood. We report here a synthetic strategy to obtain [^13^C_6_]*scyllo*-inositol and demonstrate, through isotopic tracing, its incorporation into phosphatidylinositols in different human-derived cell lines. This new stable isotopic tracer enables the investigation of the biological role of *scyllo*-inositol in mammals and beyond.

**Highlights:**

- Concise synthesis of [^13^C_6_]*scyllo*-inositol
- [^13^C_6_]*scyllo*-inositol uptake and response of endogenous inositol isomers studied in multiple cell lines
- Use of [^13^C_6_]*scyllo*-inositol as an isotopic tracer in metabolomics and lipidomics experiments
- Evidence for *scyllo*-inositol incorporation into phosphatidylinositol in mammalian cells

## Introduction

The inositol (cyclohexan-1,2,3,4,5,6-hexol) family comprises nine different stereoisomers.^1^ *Myo*-inositol is the most studied isomer, yet - with the exception of *cis*-inositol - all other members (*scyllo*-, D-*chiro*-, L-*chiro*-, *epi*-, *muco*-, *neo*- and *allo*-inositol) occur naturally as well.^2,3^ *Myo*-inositol is also the most abundant isomer in humans with plasma concentrations of approximately 30 µM, while it is found up to 5 mM in the human brain measured by magnetic resonance spectroscopy (MRS) (Figure 1A, 1B).^1,4^ This high intracellular level makes *myo*-inositol an important organic osmolyte, protecting cells from hypertonic stress especially in the brain, kidney and liver.^5^ The intracellular concentrations are controlled by the sodium *myo*-inositol co-transporters 1 and 2 (SMIT1/2), along with the proton *myo*-inositol co-transporter (HMIT).^6,7^ *Myo*-inositol also plays a major role in cellular signaling as the precursor to many secondary messengers such as inositol phosphates and inositol pyrophosphates.^8,9^ Furthermore, *myo*-inositol is an important component of phosphatidylinositols (PIs) and phosphatidylinositol phosphates (PIPs). These molecules have crucial functions in mammalian cells, for example in signal transduction, membrane dynamics and remodeling.^10^

**Figure 1:**
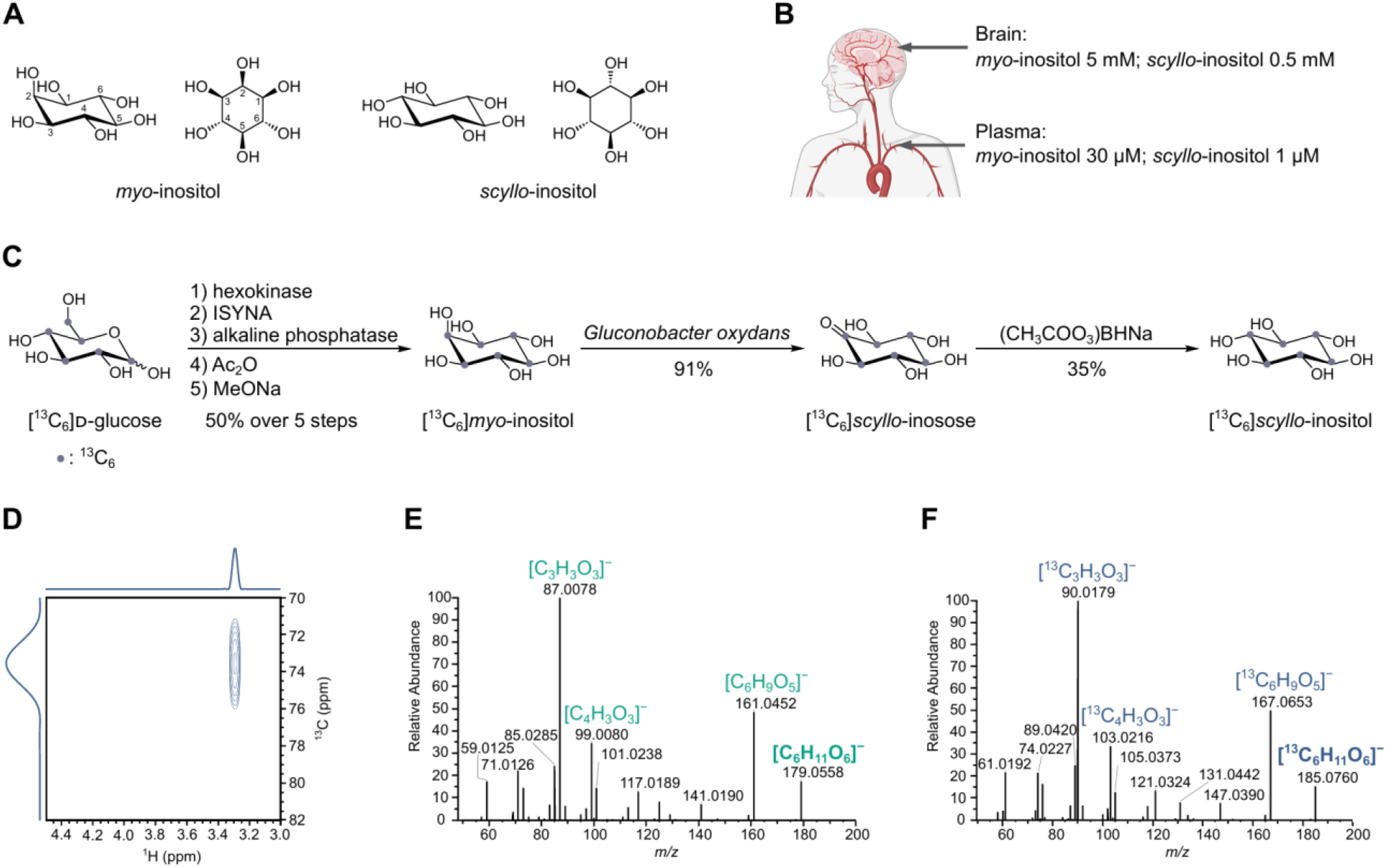
Synthesis and characterization of [^13^C_6_]*scyllo*-inositol. (A) Chemical structures of *myo*- and *scyllo*-inositol. B) Typical concentrations of *myo*- and *scyllo*-inositol in the human brain and plasma. C) Concise synthesis of [^13^C_6_]*scyllo*-inositol starting from [^13^C_6_]*myo*-inositol, which is obtained from commercial [^13^C_6_]D-glucose. The semi-enzymatic synthesis of [^13^C_6_]*myo*-inositol has been reported previously.^40,42^ ISYNA: inositol-3-phosphate synthase. D) [^1^H, ^13^C]HSQC-NMR spectrum of [^13^C_6_]*scyllo*-inositol exhibiting only a single peak due to the high symmetry of the molecule when free in solution. E–F) MS^2^ spectra for [^12^C_6_]*scyllo*-inositol (E) and [^13^C_6_]*scyllo*-inositol (F) with the respective precursor and three most abundant fragments. Some elements in 1B have been created in BioRender. Lampe, S. (2026) https://BioRender.com/gyote33.

*Scyllo*-inositol is the second most abundant inositol isomer in the human brain.^11^ It is the C2-epimer of *myo*-inositol with all hydroxyl groups in equatorial positions, which results in a fully symmetrical molecule (Figure 1A).^12^ Typical human plasma concentrations are approximately 1 µM, and cerebral *scyllo*-inositol can be investigated non-invasively by MRS because of its high concentration up to 0.5 mM (Figure 1B).^1,4,13^ These studies show that *scyllo*-inositol levels can be altered in diseases such as brain cancer, multiple sclerosis or chronic alcoholism, and are considered as prognostic biomarker.^14–18^ Histopathologically, Alzheimer’s disease (AD) can be described by the extracellular accumulation of amyloid-beta (Aβ) plaque, combined with the intracellular aggregation of hyperphosphorylated tau protein.^19^ It was shown that *scyllo*-inositol is able to stabilize small Aβ oligomers and thereby prevent aggregation.^20,21^ Therefore, *scyllo*-inositol was tested in clinical trials for the treatment of AD and was administered up to 2000 mg twice daily.^22,23^ This resulted in increased concentrations of *scyllo*-inositol in plasma, cerebrospinal fluid (CSF) and brain, but the clinical trials were discontinued because no beneficial effects could be demonstrated.^23,24^ Despite these intriguing implications for *scyllo*-inositol, it is mainly believed to act as osmolyte, and putative additional biological roles and its metabolic fate have remained unclear.^6^

To investigate the metabolic fate of a molecule of interest, isotopic tracing is a powerful method.^25^ In contrast to metabolomics experiments used for relative or absolute quantification, isotopic tracing can provide useful insights such as substrate consumption, pathway utilization or metabolic flux rewiring.^26^ It is based on the application of molecules with a different isotopic composition compared to endogenous metabolites. The tracers are enriched with an isotope that has a low natural abundance, for example, ^13^C represents approximately 1.1% of naturally occurring carbon atoms, whereas 98.9% is made up by ^12^C.^27^ The non-natural abundance of a specific isotope in the tracer allows tracking and discrimination from the endogenous metabolite by NMR, mass spectrometry (MS) or radioactive emission.^28^ Common isotopes in such tracing experiments are ^2^H, ^3^H, ^13^C, ^14^C, ^18^O, ^15^N or ^32^P.^29–32^ However, the use of certain isotopes is accompanied by undesired effects. Radioactive isotopes such as ^3^H, ^14^C or ^32^P are limited by safety measures and regulations.^33^ The radioactive decay can also limit long-term experiments because of cytotoxicity.^34^ The ^2^H isotope introduces the deuterium kinetic isotopic effect of substrates, e.g. some enzymes can discriminate between C-^1^H and C-^2^H bonds.^35^ In contrast, the stable isotopes ^13^C and ^15^N are well suited for isotopic tracing experiments as they are not harmful to cells.^36,37^ ^13^C tracers such as [^13^C_6_]glucose are commonly used and can be applied to study metabolic processes in humans.^38,39^

Our group previously established [^13^C_6_]*myo*-inositol as an isotopic tracer to study inositol phosphate metabolism in mammalian cells and phytate metabolism in the gut microbiome.^40–43^ Building on this, we now report the development of [^13^C_6_]*scyllo*-inositol for isotopic tracing. To overcome the challenges of non-existing standards for *scyllo*-inositol metabolites and isobaric analytes, LC-MS/MS and 2D-NMR were combined to investigate the metabolic fate of [^13^C_6_]*scyllo*-inositol in multiple mammalian cell lines. We demonstrate that mammalian cells can incorporate [^13^C_6_]*scyllo*-inositol into PIs, suggesting a more active metabolism of *scyllo*-inositol than previously thought.

### Design

Despite being dysregulated in various diseases and investigated in clinical trials, the biological role and metabolic fate of *scyllo*-inositol in mammalian cells remains unclear.^14–17^ To date, experiments with mammalian cells have often relied on the analysis of endogenous *scyllo*-inositol, but due to the low concentration, in particular compared to *myo*-inositol, these experiments have proven to be challenging. Isotopic tracing experiments have been conducted using [^3^H]*scyllo*-inositol to study cellular uptake in mammalian cells and its metabolism in barley seeds.^44,45^ However, the inherent radiotoxicity limited the application in long-term cellular or even *in vivo* experiments. To address the need for a stable isotopic tracer, uniformly labeled [^13^C_6_]*scyllo*-inositol was synthesized and characterized. ^13^C was chosen because it is a stable, NMR-active isotope without known cell toxicity, and it can be distinguished from the endogenous metabolite by MS.^46^ The use of [^13^C_6_]*scyllo*-inositol in isotopic tracing experiments with a combined readout of NMR and MS is an attractive approach, because it overcomes limitations of the individual analytical methods.

## Results

### Concise synthesis of [^13^C_6_]*scyllo*-inositol

A chemical synthesis of *scyllo*-inositol from *myo*-inositol, based on the oxidation of the 2-position and the selective reduction to *scyllo*-inositol, has been reported.^47^ However, this route required multiple protection and deprotection steps, which would significantly impact the overall yield. To obtain uniformly labeled [^13^C_6_]*scyllo*-inositol, we envisioned a synthesis starting from [^13^C_6_]*myo*-inositol, which can be obtained on a large scale from [^13^C_6_]glucose (Figure 1C).^40,42,48^ The microorganism *Gluconobacter oxydans* (*G.oxydans*) was utilized, whose membrane-bound dehydrogenases can selectively oxidize the 2-position of *myo*-inositol to *scyllo*-inosose (also known as *myo*-2-inosose).^49^ *G.oxydans* was cultivated in media supplemented with [^13^C_6_]*myo*-inositol for 72 h, and the supernatant was collected by centrifugation and concentrated. After precipitation and washing, pure [^13^C_6_]*scyllo*-inosose was obtained in 91% yield (Figure 1C). To obtain *scyllo*-inositol, [^13^C_6_]*scyllo*-inosose needed to be reduced. As the chemical reduction of *scyllo*-inosose resulted in a mixture of *scyllo*- and *myo*-inositol, conditions were optimized to favor [^13^C_6_]*scyllo*-inositol formation.^50^ Using sodium borohydride as reducing agent in various solvents at different temperatures did not result in a desirable ratio of [^13^C_6_]*scyllo*- to [^13^C_6_]*myo*-inositol. Switching to the milder reducing reagent sodium triacetoxyborohydride improved the ratio towards [^13^C_6_]*scyllo*-inositol (Table S1); carrying the reaction out in CH_3_CN/H_2_O (9:1) resulted in a 1:1.2 ratio of [^13^C_6_]*scyllo*-inositol to [^13^C_6_]*myo*-inositol. Separation of the two isomers was accomplished based on an adapted derivatization strategy using boric acid esterification, and [^13^C_6_]*scyllo*-inositol could be separated from derivatized [^13^C_6_]*myo*-inositol by anion exchange chromatography.^51^ Overall, [^13^C_6_]*scyllo*-inositol was obtained in 32% yield after two steps with an isotopic purity of 99% (Figure 1C). In the ^1^H and the ^13^C NMR spectra, [^13^C_6_]*scyllo*-inositol exhibited a singlet, illustrating that *scyllo*-inositol is a fully symmetrical molecule. For the same reason, a [^1^H, ^13^C] heteronuclear single quantum correlation (HSQC)-NMR experiment, which selectively detected ^13^CH groups, displayed only a single peak (Figure 1D). If the symmetry of the molecule would be broken, a characteristic triplet pattern due to carbon-carbon coupling would be visible in the C dimension, which could facilitate the identification of [^13^C_6_]metabolites. A MS^2^ spectrum of [^12^C_6_]*scyllo*-inositol showed the precursor and three most abundant fragments that had also been reported for [^12^C_6_]*myo*-inositol (Figure 1E, Figure S1A).^1^ [^13^C_6_]*scyllo*-inositol exhibited the same fragmentation pattern as the [^12^C_6_]isotopologue (Figure 1F, Figure S1B). The precursor and fragments displayed the expected *m/z* shift corresponding to the mass difference between ^12^C and ^13^C.

### *Myo*-inositol depleted mammalian cells can be cultivated in [^13^C_6_]*scyllo*-inositol

With [^13^C_6_]*scyllo*-inositol in hand, we wanted to investigate its metabolic fate in mammalian cells. To identify suitable time points for such experiments, it was necessary to determine whether [^13^C_6_]*scyllo*-inositol reached a steady-state and if it behaved similarly to *myo*-inositol in general. The preferred model system was a neural tissue-derived cell line, because high concentrations of *scyllo*-inositol have been reported in the human brain and aberrant *scyllo*-inositol levels have been linked to various diseases, including brain cancer.^4,18,52,53^ Therefore A172 cells were chosen, a glioblastoma derived cell line with reportedly high expression of the inositol transporter HMIT among glioblastomas.^54,55^

First, the proliferation of A172 cells was tested in medium with defined concentrations of inositol isomers and dialyzed fetal bovine serum (FBS) (Table S2).^56^ A172 cells were cultured in DMEM containing either 100 µM [^12^C_6_]*myo*-inositol, 100 µM [^12^C_6_]*scyllo*-inositol, or a combination of 90 µM [^12^C_6_]*myo*- and 10 µM [^12^C_6_]*scyllo*-inositol, which resembles cerebral concentrations. Proliferation was monitored for four days by a water-soluble tetrazolium (WST) assay and compared to A172 cells cultured in commercial DMEM with standard FBS. No notable differences between the distinct conditions were observed, indicating that A172 cells were suited as a model system to study *scyllo*-inositol metabolism (Figure 2A).

**Figure 2:**
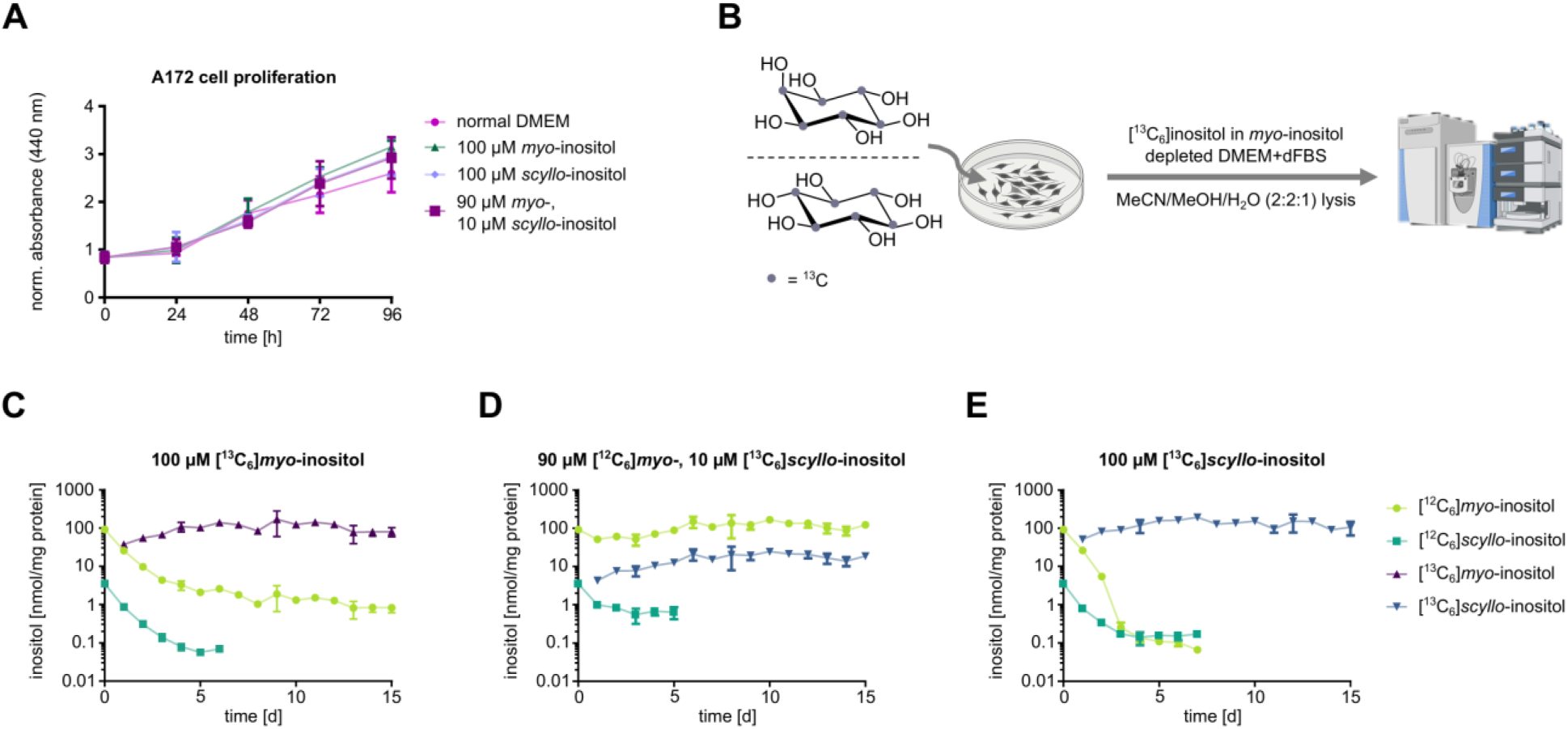
Import of [^13^C_6_]*scyllo*-inositol in A172 cells. A) Cell proliferation of A172 cells cultivated in defined inositol concentrations in cell media supplemented with dFBS. Proliferation was measured by a WST assay. B) Schematic representation of long-term experiments to study the import of [^13^C_6_] inositol isomers and the response of endogenous [^12^C_6_] inositol isomers in A172 cells. C–E) Time course of intracellular inositol levels in A172 cells cultivated in 100 µM [^13^C_6_]*myo*-inositol (C) the combination of 90 µM [^12^C_6_]*myo*-inositol and 10 µM [^13^C_6_]*scyllo*-inositol (D) or 100 µM [^13^C_6_]*scyllo*-inositol (E). Inositol levels were measured by HILIC-MS/MS and normalized to protein concentration. All data is shown as mean ± standard deviation (n = 3 biological replicates), except in C) treatment with 100 µM [^13^C_6_]*myo*-inositol at day 14. This data point is shown as mean ± standard deviation (n = 2 biological replicates). Some elements in 2B have been created in BioRender. Lampe, S. (2026) https://BioRender.com/gyote33.

Next, we investigated how the concentrations of cellular inositol isomers were affected when A172 cells were cultured in DMEM with defined concentrations of inositol isomers. A172 cells were cultured up to 15 days, and the concentrations of intracellular [^12^C_6_]/[^13^C_6_]*myo*- and [^12^C_6_]/[^13^C_6_]*scyllo*-inositol were measured by an adapted hydrophilic liquid interaction (HILIC)-MS/MS method using a mild lysis protocol that did not release covalently bound inositol (Figure 2B).^1,57^ Cultivation in 100 µM [^13^C_6_]*myo*-inositol resulted in a fast uptake and a steady state between [^13^C_6_]*myo-* and [^12^C_6_]*myo-*inositol after five days. [^13^C_6_]*myo*-inositol accounted for approximately 95% of the total intracellular *myo*-inositol concentration. Under these conditions, intracellular [^12^C_6_]*scyllo*-inositol levels slowly decreased, and the concentration reached the limit of detection after six days (Figure 2C). Using 90 µM [^12^C_6_]*myo*- and 10 µM [^13^C_6_]*scyllo*-inositol did not affect the intracellular [^12^C_6_]*myo*-inositol concentration and resulted in an increase of [^13^C_6_]*scyllo*-inositol, which reached a steady state after two days. A172 cells rapidly equilibrated the cellular concentrations of [^12^C_6_]*myo*-inositol and [^13^C_6_]s*cyllo*-inositol to reflect the 9:1 ratio in the culture medium (Figure 2D). Cultivation of A172 cells in 100 µM [^13^C_6_]*scyllo*-inositol resulted in a steady state of intracellular [^13^C_6_]*scyllo*-inositol after five days, and the concentration was similar to the basal [^12^C_6_]*myo*-inositol concentration. In contrast to the other conditions, a rapid decline of intracellular [^12^C_6_]*myo*-inositol was observed. Within four days, the concentration decreased by more than three orders of magnitude and later fell below the limit of detection. Also, the concentration of [^12^C_6_]*scyllo*-inositol showed a fast decrease and was below the limit of detection after seven days (Figure 2E). These experiments demonstrated that A172 cells can be cultivated in 100 µM [^13^C_6_]*scyllo*-inositol, which is readily imported, while endogenous pools of intracellular [^12^C_6_]*myo*- and [^12^C_6_]*scyllo*-inositol were depleted.

### A172 cells incorporate *scyllo*-inositol into phosphatidylinositols

Having confirmed the import of [^13^C_6_]*scyllo*-inositol into A172 cells, we were curious whether *scyllo*-inositol has a similar metabolic fate as *myo*-inositol, which gets incorporated into PIs.^58^ The incorporation of *scyllo*-inositol into PIs had only been observed previously in aleurone cells of barley seeds and the unicellular eukaryotic model organism *Tetrahymena vorax* using radiolabeling or GC-MS analysis of *scyllo*-inositol after PI hydrolysis.^44,59–62^ However, to our knowledge, no data exists for mammalian cells. A172 cells were cultivated in 100 µM [^13^C_6_]*scyllo*-inositol for eight days, as our time-course experiments indicated a sufficient exchange of endogenous intracellular inositol isomers. Three independent approaches were then used to probe for PI incorporation, two of them were MS-based (Figure 3A).

**Figure 3:**
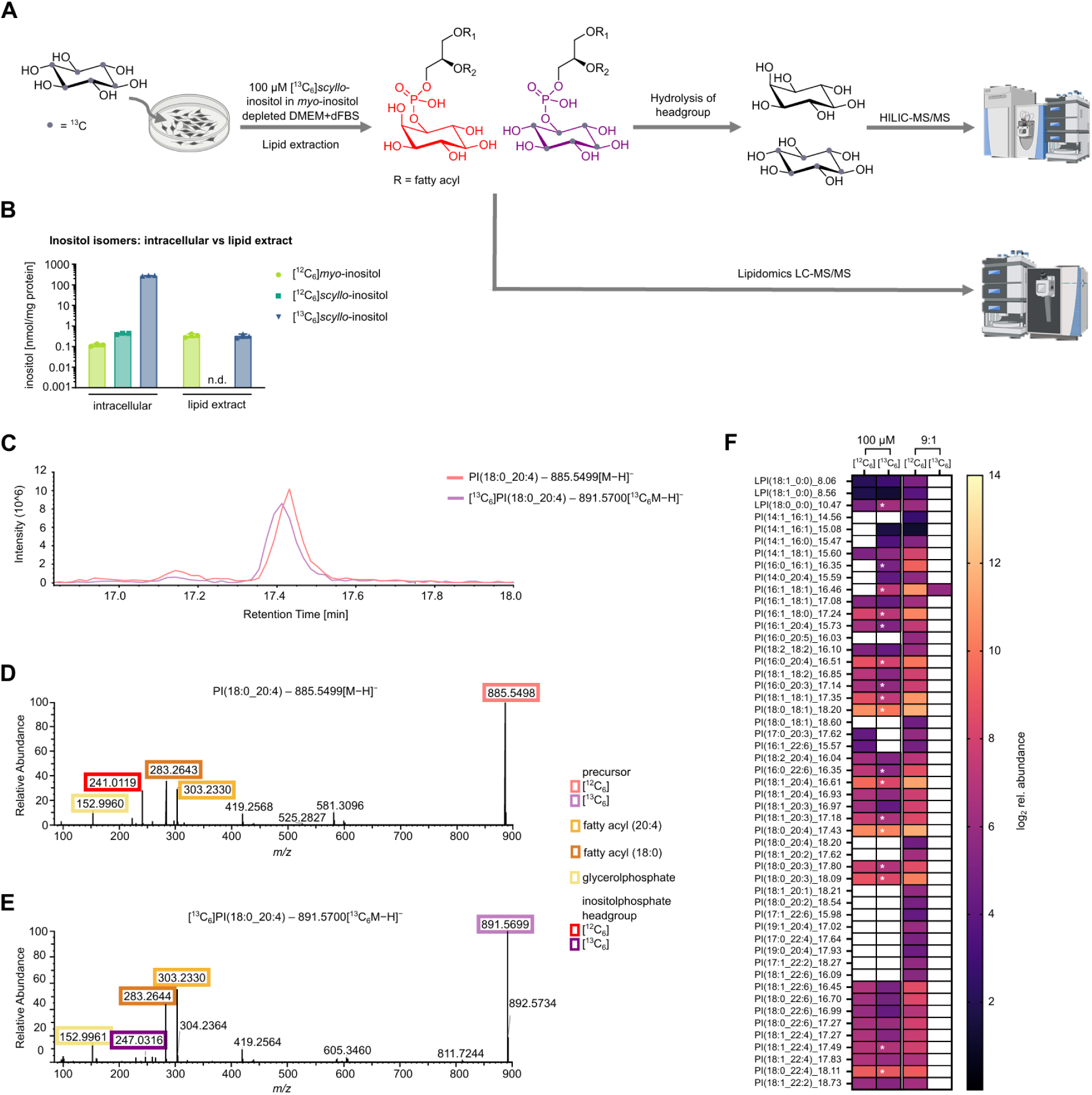
[^13^C_6_]Inositol incorporation into PIs. A) Schematic workflow for labeling of A172 cells followed by lipid extraction and headgroup analysis or LC-MS/MS based lipidomics. B) A172 cells were cultivated in 100 µM [^13^C_6_]*scyllo*-inositol for eight days and inositol isomers were analyzed by HILIC-MS/MS. The data is shown as mean ± standard deviation (n = 3 biological replicates). n.d.: not determined. C) Extracted ion chromatograms of PI(18:0_20:4) with co-elution of [^12^C_6_] and [^13^C_6_] precursor. D–E) MS^2^ spectra PI(18:0_20:4) with precursor and fragments of headgroup, glycerolphosphate and both fatty acyls for [^12^C_6_] (D) and [^13^C_6_] species (E). F) Heatmap representing the relative abundance of all putative and identified lyso-PI (LPI) and PI species ([^12^C_6_] and [^13^C_6_]) in A172 cells cultivated for eight days in either 100 µM [^13^C_6_]*scyllo*-inositol or the combination of 90 µM [^12^C_6_]*myo*-inositol and 10 µM [^13^C_6_]*scyllo*-inositol (9:1), respectively. Identified [^13^C_6_]PI species are marked with a white asterisk. Data is shown in log_2_ values (n = 3 biological replicates). LPIs and PIs are annotated on the fatty acyl level and corresponding retention times are indicated in min. Some elements in 3A have been created in BioRender. Lampe, S. (2026) https://BioRender.com/gyote33.

In a first approach, the headgroups were analyzed following PI extraction. Lipids were extracted from A172 cell pellets using chloroform (CHCl_3_) and subsequently hydrolyzed with perchloric acid.^63^ After neutralization, the supernatant was analyzed for [^12^C_6_]/[^13^C_6_]inositol isomers by HILIC-MS/MS. The extraction protocol was extended by additional washing steps with ddH_2_O to prevent carry-over of intracellular inositol isomers, which are not lipid-derived (Figure S2A, B). Analysis of cell samples cultivated in 100 µM [^13^C_6_]*scyllo*-inositol for eight days uncovered approximately equal concentrations of [^13^C_6_]*scyllo*-inositol (0.3 nmol/mg protein) and [^12^C_6_]*myo*-inositol (0.3 nmol/mg protein), while [^12^C_6_]*scyllo*-inositol and [^13^C_6_]*myo*-inositol were not detected (Figure 3B). Levels of extracted and intracellular [^12^C_6_]*myo*-inositol had the same order of magnitude, whereas intracellular levels of [^13^C_6_]*scyllo*-inositol were approximately 1000-fold higher than extracted [^13^C_6_]*scyllo*-inositol (Figure 3B). The combined amount of [^13^C_6_]*scyllo*-inositol and [^12^C_6_]*myo*-inositol extracted from lipids was in the same range as literature reports on PI concentrations in cultured mouse cortical neurons and in post mortem samples from human brains.^58,64^ For the combination of 90 µM [^12^C_6_]*myo*- and 10 µM [^13^C_6_]*scyllo*-inositol, small amounts of [^13^C_6_]*scyllo*-inositol were also detected in the chloroform extract, although approximately five times the amount of cellular material was needed (Figure S2C). The HILIC-MS/MS data indicated that [^13^C_6_]*scyllo*-inositol could indeed be incorporated into lipids as a headgroup, presumably as PIs.

To further confirm the putative incorporation of [^13^C_6_]*scyllo*-inositol into PI species, lipidomics was used as a second approach to directly detect labeled lipid species. A172 cells were cultivated in 100 µM [^13^C_6_]*scyllo*-inositol or the combination of 90 µM [^12^C_6_]*myo*- and 10 µM [^13^C_6_]*scyllo*-inositol for eight days, and PIs were extracted using methyl *tert*-butyl ether (MTBE) (Figure 3A).^65^ After untargeted LC-MS/MS measurements, the [^12^C_6_]PI species were assigned using Lipostar2 software.^66^ Next, two MS methods with targeted data-dependent acquisition (t-DDA) mode were generated. The first method included all assigned [^12^C_6_]PI species and their corresponding [^13^C_6_]PI species. The second method only included the [^13^C_6_]PI species, to increase sensitivity. The samples were measured with both methods, and the first t-DDA data were used for relative quantification, while t-DDA data from the second method were used for identification. Peaks were assigned as putative [^13^C_6_]PI species if they co-eluted with their corresponding [^12^C_6_]PI species without sufficient spectra for the identification as [^13^C_6_]PI. Exemplified by the extracted ion chromatograms of the most abundant PI species (18:0_20:4), both precursors showed the exact mass difference of [^13^C_6_] to [^12^C_6_] (Figure 3C). A specific [^13^C_6_]PI species was identified, if the following characteristic fragments were detected in the MS^2^ spectra: the glycerolphosphate backbone, the [^13^C_6_]inositol-headgroup and the respective fatty acyls. As illustrated by the spectra for the PI(18:0_20:4) species, only the inositol-headgroup fragments differed between the [^13^C_6_]PI and the [^12^C_6_]PI isomer (Figure 3D, 3E). By applying the aforementioned criteria, 17 distinct PI species with a [^13^C_6_]headgroup were identified in A172 cells cultivated in 100 µM [^13^C_6_]*scyllo*-inositol, while an additional 19 could only be assigned as putative [^13^C_6_]PI species. For the A172 cells cultured in the combination of 90 µM [^12^C_6_]*myo*- and 10 µM [^13^C_6_]*scyllo*-inositol, only one PI species could be putatively assigned with a [^13^C_6_]headgroup (Figure 3F). These data confirmed the incorporation of [^13^C_6_]inositol isomers into PI species in A172 cells. While this method allowed for their identification, it did not yield further structural information about the respective [^13^C_6_]inositol headgroup.

To confirm the structure of the headgroup, 2D NMR spectroscopy was applied as a third approach. A172 cells were cultivated in 100 µM [^13^C_6_]*scyllo*-inositol for eight days and extracted with MTBE, as was done for the lipidomics samples (Figure 4A). The extracted PIs were dissolved in CDCl_3_/CD_3_OD (4:1) and analyzed by [^1^H, ^13^C] HSQC-NMR using a BIRD filter.^67^ The NMR spectrum showed three signals with chemical shifts anticipated for inositol C-H groups, in both ^1^H and ^13^C dimensions (Figure 4B, Figure S3A). The characteristic triplet pattern in the C dimension arose from coupling of ^13^C nuclei to neighboring ^13^C nuclei. These signals were absent in the corresponding [^12^C_6_]*scyllo*-inositol control sample (Figure 4B, Figure S3A). The signals at ^1^H: 3.88 ppm/^13^C: 80.02 ppm (C1 H), ^1^H: 3.27 ppm/^13^C: 73.77 ppm (C3, 4, 5 H) and ^1^H: 3.41 ppm/^13^C: 73.31 ppm (C2, 6 H) had relative integrals of 1.0:2.4:1.5. Considering the complex nature of the sample, and the fact that HSQC spectra are not fully quantitative, the ratio was approximately 1:3:2, which fits to inositol isomers. For comparison, a [^1^H, ^13^C]HSQC-NMR spectrum of A172 cells cultivated in 100 µM [^13^C_6_]*myo*-inositol was measured after MTBE extraction. The spectrum displayed six signals that were distinct from those derived from [^13^C_6_]*scyllo*-inositol treated A172 cells (Figure 4B, Figure S3B, C). A [^1^H, ^13^C]HSQC combined with a clean in-phase correlation spectroscopy (CLIP-COSY) spectrum was used to assign the signals in the [^13^C_6_]*myo*-inositol sample (Figure S3D). The cross-correlations of the signals, and comparison with a literature spectrum, allowed us to assign the signals to *myo*-inositol-PI.^68^ The [^1^H, ^13^C]HSQC-CLIP-COSY spectrum of the [^13^C_6_]*scyllo*-inositol sample displayed cross-correlations between the signal at ^1^H: 3.88 ppm/^13^C: 80.02 ppm and ^1^H: 3.41 ppm/^13^C: 73.31 ppm (Figure 4C). Given their relative integrals, the signal at ^1^H: 3.41 ppm/^13^C: 73.31 ppm was indicative of two protons adjacent to the signal at ^1^H: 3.88 ppm/^13^C: 80.02 ppm. The signal at ^1^H: 3.27 ppm/^13^C: 73.77 ppm was not fully resolved in the [^1^H, ^13^C]HSQC-CLIP-COSY spectrum and corresponds to three protons. A weak cross-correlation between the smaller lower part of the signal at ^1^H: 3.27 ppm/^13^C: 73.77 ppm and the signal at ^1^H: 3.41 ppm/^13^C: 73.31 ppm was observed. The spectra of MTBE extracts from A172 cells cultivated in [^13^C_6_]*scyllo*-inositol indicated the presence of a derivatized [^13^C_6_]*scyllo*-inositol isomer. Comparison with the respective [^13^C_6_]*myo*-inositol sample suggested derivatization at one position. The number of signals in the [^13^C_6_]*scyllo*-inositol sample and their respective integrals match the symmetry of [^13^C_6_]*scyllo*-inositol derivatized at one position.^69^ The non-availability of a *scyllo*-inositol-PI standard is a challenge, because the inositol isomer cannot be unambiguously assigned as *scyllo*-inositol. However, the data of the hydrolyzed samples from the same experiment were also considered in the interpretation. Thus, we are confident that the isolated [^13^C_6_]species is *scyllo*-PI. Thereby, we showcased that the combination of multiple analytical methods can overcome such a challenge and confirm the incorporation of [^13^C_6_]*scyllo*-inositol into PIs in A172 cells after being cultured for eight days in medium containing 100 µM [^13^C_6_]*scyllo*-inositol.

**Figure 4:**
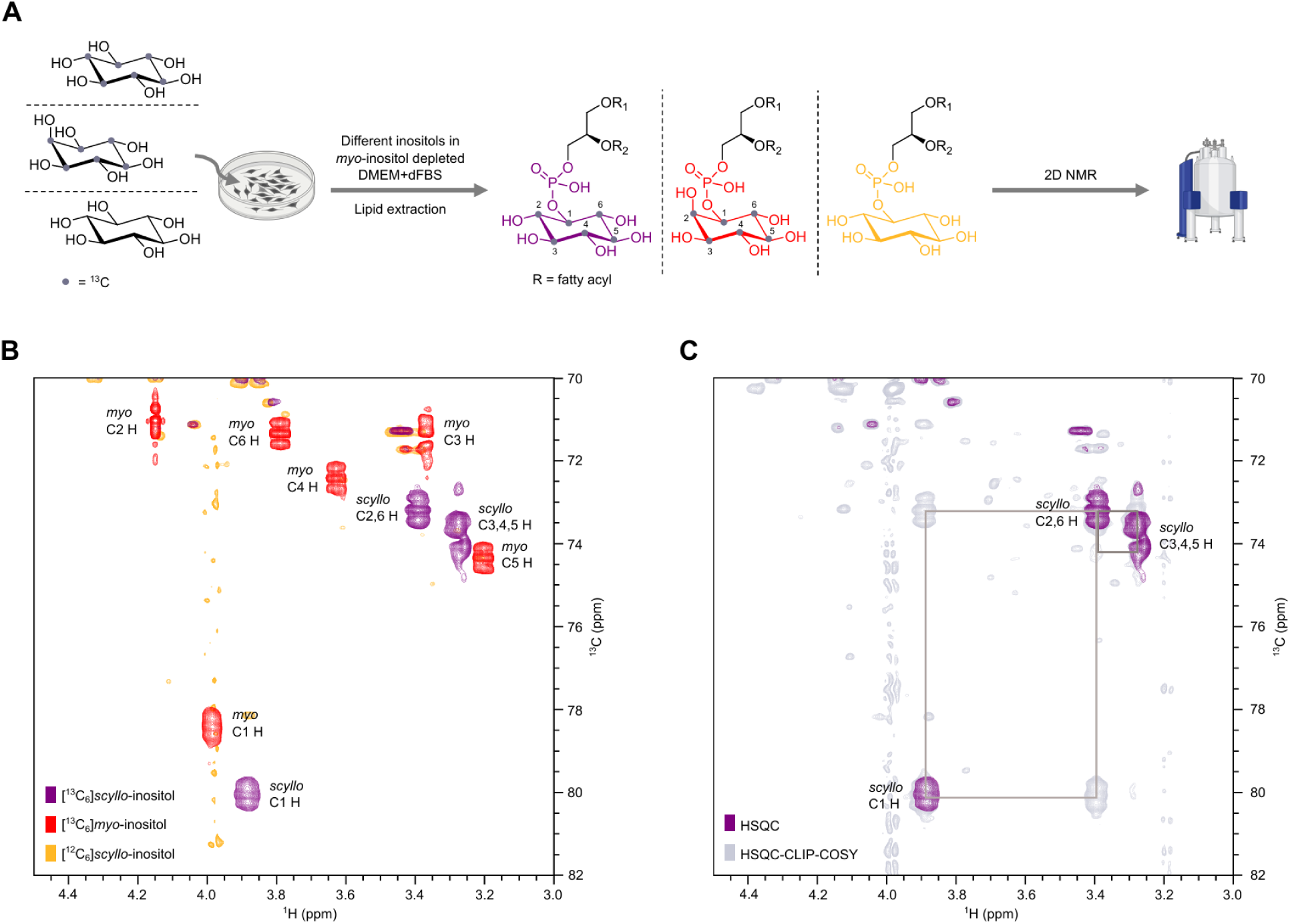
Confirmation of [^13^C_6_]*scyllo*-inositol incorporation into lipids. A) Schematic workflow for labeling of A172 cells followed by lipid extraction and direct headgroup analysis by 2D NMR. B) Overlay of [^1^H, ^13^C]HSQC-NMR spectra using a BIRD filter from lysate of A172 cells cultivated in either 100 µM [^13^C_6_]*scyllo*-inositol, 100 µM [^13^C_6_]*myo*-inositol or 100 µM [^12^C_6_]*scyllo*-inositol. C) Overlay of [^1^H, ^13^C]HSQC and HSQC-CLIP-COSY spectra to highlight cross-correlation in lysate of A172 cells cultivated in 100 µM [^13^C_6_]*scyllo*-inositol. Some elements in 4A have been created in BioRender. Lampe, S. (2026) https://BioRender.com/gyote33.

### *Scyllo*-inositol is incorporated into phosphatidylinositols in HEK293T and HCT116 cells,mm

The incorporation of [^13^C_6_]*scyllo*-inositol into PIs in A172 cells prompted us to investigate other mammalian cell lines. HEK293T and HCT116 cells have been extensively used to study inositol isomers and their metabolism.^40,45,57,70,71^ Both cell lines proliferated well in DMEM supplemented with dialyzed FBS containing 100 µM [^13^C_6_]*scyllo*-inositol (Figure S4). HEK293T and HCT116 cells were then cultivated for eight or ten days, respectively, and the inositol metabolism was analyzed by different methods. First, the concentrations of intracellular [^12^C_6_]/[^13^C_6_]*myo*- and [^12^C_6_]/[^13^C_6_]*scyllo*-inositol were measured by HILIC-MS/MS. HEK293T cells behaved similar to A172 cells and showed an uptake of [^13^C_6_]*scyllo*-inositol (100 nmol/mg protein) while [^12^C_6_]*myo*-inositol (0.1 nmol/mg protein) and [^12^C_6_]*scyllo*-inositol (0.1 nmol/mg protein) were depleted (Figure 5A). While the concentration of [^13^C_6_]*scyllo*-inositol was lower compared to A172 cells, the ratio of [^13^C_6_]*scyllo*-inositol to [^12^C_6_]*myo*-inositol was very similar in both cell lines. Interestingly, HCT116 cells showed a much lower uptake of [^13^C_6_]*scyllo*-inositol (20 nmol/mg protein) than A172 and HEK293T cells. In contrast, the concentration of [^12^C_6_]*myo*-inositol (0.9 nmol/mg protein) was much higher compared to A172 and HEK293T cells, which could indicate a different inositol metabolism (Figure 5A).

**Figure 5:**
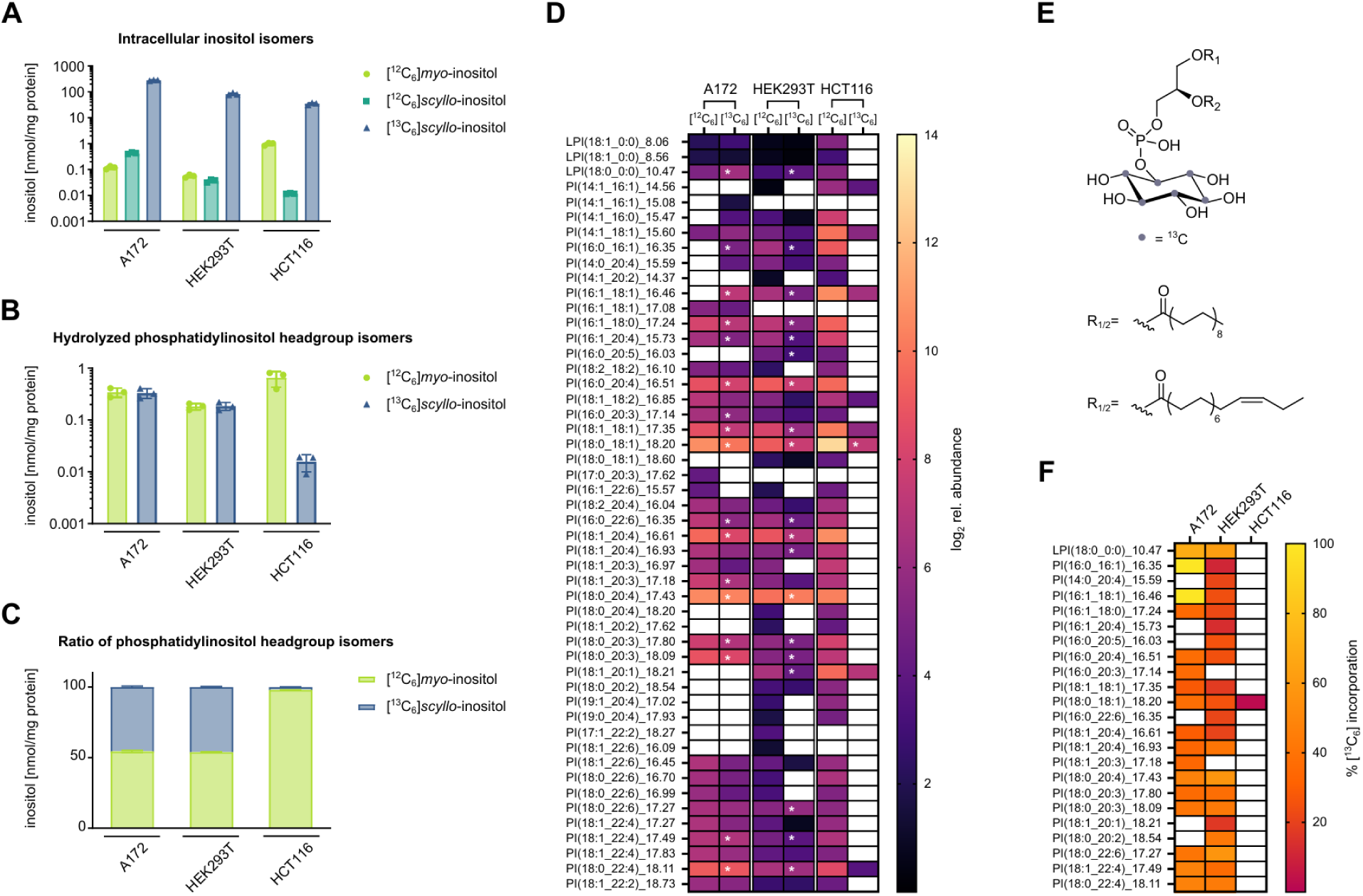
Confirmation of [^13^C_6_]*scyllo*-inositol incorporation in multiple mammalian cell lines. A–B) Cells were cultivated for eight days in 100 µM [^13^C_6_]*scyllo*-inositol. Intracellular inositol isomers were analyzed by HILIC-MS/MS (A) or after lipid extraction and subsequent hydrolysis (B). C) Ratio of [^12^C_6_]*myo*-inositol to [^13^C_6_]*scyllo*-inositol found after lipid extraction and subsequent hydrolysis. The data is shown as mean ± standard deviation (n = 3 biological replicates). D) Heatmap representing the relative abundance of all putative and identified PI species ([^12^C_6_] and [^13^C_6_]) in A172, HEK293T and HCT116 cells cultivated for eight days in 100 µM [^13^C_6_]*scyllo*-inositol. Identified [^13^C]PI species are marked with a white asterisk. Data is expressed in log_2_ values (n = 3 biological replicates). E) Chemical structure of PI(18:0_18:1), which showed incorporation of [^13^C_6_]*scyllo*-inositol in all cell lines. F) Heatmap representing the average incorporation of [^13^C_6_]*scyllo*-inositol into the identified PI species (n = 3 biological replicates).

Next, CHCl_3_ lipid extractions from HEK293T and HCT116 cells were performed, followed by perchloric acid hydrolysis of the headgroups, and [^12^C_6_]/[^13^C_6_]inositol isomers were measured by HILIC-MS/MS. Similar to A172 cells, HEK293T cells showed an approximate 1:1 ratio of [^13^C_6_]*scyllo*-inositol 0.2 nmol/mg protein) to [^12^C_6_]*myo*-inositol (0.2 nmol/mg protein), suggesting that [^13^C_6_]*scyllo*-inositol could be incorporated into lipids as a headgroup (Figure 5B). By contrast, HCT116 cells showed a 1:99 ratio of [^13^C_6_]*scyllo*-inositol (0.01 nmol/mg protein) to [^12^C_6_]*myo*-inositol (0.9 nmol/mg protein), indicating only minimal incorporation into lipids (Figure 5C). Lastly, MS-based lipidomics of HEK293T and HCT116 cells cultivated in 100 µM [^13^C_6_]*scyllo*-inositol were conducted to test for PI incorporation. In HEK293T cells, 34 [^13^C_6_]PI species were observed, and by using the criteria as for A172 cells, 22 [^13^C_6_]PI species could be confirmed, while 12 PI species remained putatively assigned (Figure 5D). Together with the HILIC-MS/MS data after CHCl_3_ extraction, the lipidomics data confirmed that HEK293T cells also incorporate [^13^C_6_]*scyllo*-inositol into PIs. Only one [^13^C_6_]PI species was identified in HCT116 cells, and seven [^13^C_6_]PI species were putatively assigned, further indicating cell-line specific differences in inositol metabolism. The [^13^C_6_]PI species (18:0_18:1) was identified in all three cell lines (Figure 5E).

t-DDA data of the [^12^C_6_]/[^13^C_6_] method were used to calculate the incorporation of the [^13^C_6_]inositol headgroup in all cell lines (Figure 5F). The average incorporation across all identified [^13^C_6_]PI species was approximately 46% in A172 cells and two PI species, 16:0_16:1 and 16:1_18:1, showed 100% incorporation. HEK293T cells showed a similar average incorporation of approximately 31%. The [^13^C_6_]PI species identified in HCT116 cells had an incorporation of approximately 2%, consistent with much lower levels of [^13^C_6_]*scyllo*-inositol measured in CHCl_3_ extraction after hydrolysis.

## Discussion

Isotopic tracing is a powerful method for the investigation of the metabolic fate of molecules. In this work, we synthesized [^13^C_6_]*scyllo*-inositol as a new tool and demonstrated in a proof-of-principle study its applicability in cellular models by studying cellular uptake and its incorporation into PIs in A172, HEK293T and HCT116 cells.

Our synthetic strategy has minimal requirements, because *G.oxydans* is a wild-type organism and no bioreactor is needed for its cultivation.^49^ The reduction of [^13^C_6_]*scyllo*-inosose is also straightforward, since no inert atmosphere or dry conditions are needed for this robust step. By optimizing the reaction conditions, nearly a 1:1 ratio of [^13^C_6_]*scyllo*-inositol and [^13^C_6_]*myo*-inositol can be obtained. Separation of both isomers by anion exchange chromatography provides pure [^13^C_6_]*scyllo*-inositol and offers the option to recover [^13^C_6_]*myo*-inositol.^51^

In contrast to a previous report about HEK293T cells cultivated in 500 µM *scyllo*-inositol, cultivation in 100 µM *scyllo*-inositol and dialyzed FBS was well tolerated by A172, HEK293T and HCT116 cells.^71^ [^13^C_6_]*scyllo*-inositol is imported rapidly by A172 cells and reaches intracellular concentrations similar to [^12^C_6_]*myo*-inositol. These levels were stable for 14 days, indicating that A172 cells can replace [^12^C_6_]*myo*-inositol with [^13^C_6_]*scyllo*-inositol, potentially to take over the osmotic function.^6^ Cultivation in 100 µM [^13^C_6_]*scyllo*-inositol led to a strong and rapid decline of [^12^C_6_]*myo*-inositol, while 100 µM [^13^C_6_]*myo*-inositol resulted in a steady-state between [^12^C_6_] and [^13^C_6_]*myo*-inositol. Possible reasons for the decline of [^12^C_6_]*myo*-inositol could be efflux or consumption in metabolic processes, for which it could not be substituted with [^13^C_6_]*scyllo*-inositol. These findings are consistent with observations from clinical trials with *scyllo*-inositol, in which measurements by MRS showed a decrease of *myo*-inositol concurrent with an increase of *scyllo*-inositol.^72^ [^13^C_6_]*scyllo*-inositol now enables pulse-chase experiments with labeled molecules such as [^13^C_2_]*myo*-inositol and [^13^C_3_]glucose to further investigate the rapid depletion of [^12^C_6_]*myo*-inositol when A172 cells were cultivated in 100 µM [^13^C_6_]*scyllo*-inositol.^42,57,73^ When cultivated in a combination of 90 µM [^12^C_6_]*myo*-inositol and 10 µM [^13^C_6_]*scyllo*-inositol, this ratio is reflected by intracellular concentrations of both isomers in A172 cells. This observation could suggest that A172 cells do not discriminate between both isomers and import unselectively from the medium, as already shown in HEK293T cells using isotopic tracing.^45^ HEK293T and HCT116 cells also import [^13^C_6_]*scyllo*-inositol in the absence of [^12^C_6_]*myo*-inositol. Both cell lines had lower concentrations of intracellular [^13^C_6_]*scyllo*-inositol, which is in agreement with overall lower concentrations of intracellular *myo*-inositol reported compared to A172 cells.^57^ HCT116 cells have the lowest concentration of [^13^C_6_]*scyllo*-inositol and sustained an approximately ten times higher concentration of [^12^C_6_]*myo*-inositol. A possible explanation could be that HCT116 cells rather rely more strongly on the *de novo* synthesis of *myo*-inositol from glucose than direct import compared to other cell lines.^74^

The metabolic fate of *scyllo*-inositol is unknown in mammalian cells. However, for neurological disorders, significant changes in *scyllo*-inositol and dysregulation of PI metabolism have been reported in the human brain.^14–17,75,76^ Because *myo*-inositol is incorporated into PIs, we were curious whether *scyllo*-inositol can also be incorporated into PIs in mammalian cells, as was already shown for aleurone cells of barley seeds and *Tetrahymena vorax*.^44,59–62^ Isotopic tracing with [^13^C_6_]*scyllo*-inositol was used to test for PI incorporation in A172, HEK293T and HCT116 cells. As there was no single experiment to prove the existence of [^13^C_6_]*scyllo*-PI, we combined three different methods. First, an organic extraction of lipids followed by acidic hydrolysis was used to release the inositol headgroup. HILIC-MS/MS measurements showed a 1:1 ratio [^12^C_6_]*myo*-inositol to [^13^C_6_]*scyllo*-inositol for A172 and HEK293T cells, whereas a 99:1 ratio was detected in HCT116 cells. This indirect method indicated that [^13^C_6_]*scyllo*-inositol could be incorporated into lipids, although the high concentrations of intracellular [^13^C_6_]*scyllo*-inositol could cause a carry-over. Lipidomics based on LC-MS/MS was applied to investigate intact lipid species and identify the respective species with a [^13^C_6_]headgroup. The measurements identified PI species with a [^13^C_6_] mass shift compared to the respective [^12^C_6_]PI species. While several [^13^C_6_]PI species were unambiguously identified in A172, HEK293T and HCT116 cells, this method lacks chromatographic separation of *myo*- and *scyllo*-PI, preventing the differentiation of the headgroups. We therefore took advantage of the good detectability of [^13^C_6_]*scyllo*-inositol by NMR and analyzed the lipid extracts from A172 cells by 2D NMR. Although this experiment provided no information about the PI species, it confirmed that the [^13^C_6_] headgroup is indeed *scyllo*-inositol. The combination of LC-MS/MS based identification of [^13^C_6_]PI species and the NMR-based confirmation of a [^13^C_6_]*scyllo*-inositol headgroup provide evidence that we identified [^13^C_6_]*scyllo*-PI in mammalian cells. This proof-of-concept study of [^13^C_6_]*scyllo*-inositol as an isotopic tracer addressed the long-standing question about the metabolic fate of *scyllo*-inositol. As there are no standards for *scyllo*-inositol metabolites available, we demonstrated that such challenges can be overcome by using multiple analytical methods to characterize [^13^C_6_]*scyllo*-inositol and its metabolites.

The number of identified [^13^C_6_]*scyllo*-PI species was similar in A172 and HEK293T cells, while only one species could be identified in HCT116 cells. A172 cells showed the highest average incorporation, which could be due to higher intracellular [^13^C_6_]*scyllo*-inositol concentrations. During PI biosynthesis, the inositol headgroup is transferred onto cytidine diphosphate (CDP)-diacylglycerol (DG) by CPD-DG-inositol transferase. The canonical substrate is *myo*-inositol, but for barley aleurone cells, it was shown that *scyllo*-inositol can also serve as substrate, suggesting no differentiation of inositol isomers.^59^ PI(18:0_20:4 or 38:4) was the most abundant species in A172 and HEK293T cells, which had 50% [^13^C_6_]*scyllo*-inositol incorporation in both cell lines. This data is in agreement with literature findings that PI(18:0_20:4 or 38:4) is the most abundant PI species in mammalian cells and tissues.^77–79^ The only [^13^C_6_]PI species identified in HCT116 was PI(18:0_18:1 or 36:1) with a very low incorporation of [^13^C_6_]*scyllo*-inositol, and this species was also identified in A172 and HEK293T cells. A fast PI turnover has been shown for PI(36:1) with a high dependency on *de novo* synthesis, which could explain why PI(36:1) was identified as [^13^C_6_]PI species in A172, HEK293T and HCT116 cells.^80,81^ The overall low incorporation of [^13^C_6_]*scyllo*-inositol in HCT116 could originate from PI synthesis being more dependent on *de novo* synthesis of *myo*-inositol from glucose or a more active recycling pathway for PI.^74,77^

The development of [^13^C_6_]*scyllo*-inositol allowed previously non-feasible isotopic tracing experiments and the in-depth investigation of *scyllo*-inositol metabolism. The results revealed an unknown metabolic pathway of *scyllo*-inositol in mammalian cells, in addition to being an osmolyte.^6^ Although depletion of [^12^C_6_]*myo*-inositol and supplementation of [^13^C_6_]*scyllo*-inositol is non-physiological, 100 µM [^13^C_6_]*scyllo*-inositol is in the same concentration range as CSF concentrations measured in clinical trials after eleven days of 2000 mg administration twice daily.^24^ Together with the observed decrease of *myo*-inositol in these trials, our culture concentrations resemble realistic conditions in the human brain, which makes the observed incorporation into PIs even more interesting.^72^ Cultivation in a combination of 90 µM [^12^C_6_]*myo*-inositol and 10 µM [^13^C_6_]*scyllo*-inositol reflects the physiological concentration and ratio of both inositol isomers in human CSF.^82^ Under these conditions, only one putative PI was identified, which fits much lower concentrations of intracellular [^13^C_6_]*scyllo*-inositol. For investigations of cells cultivated under physiological inositol conditions, longer cultivation times, a higher cell number, and lipidomics methods with increased sensitivity could provide more insights.

Cerebral *scyllo*-inositol levels are elevated in diseases such as brain cancer, multiple sclerosis or chronic alcoholism, but the underlying mechanism are unclear.^14–17^ Dysregulations in import or metabolism are discussed, and a potential role of an altered gut microbiome metabolism is hypothesized.^83^ The development of [^13^C_6_]*scyllo*-inositol as an isotopic tracer and first data of its incorporation into PIs will open up new possibilities to investigate these open questions. In addition to cell models, [^13^C_6_]*scyllo*-inositol can be used in *ex vivo*/*in vivo* models or humans. The NMR activity of the tracer also enables live studies in relevant disease models using advanced techniques such as high-resolution magic angle spin NMR.^84^

In conclusion, we developed [^13^C_6_]*scyllo*-inositol as an isotopic tracer and provided evidence that *scyllo*-inositol is incorporated into PIs in mammalian cells. We also uncovered cell-line specific differences in incorporation, which hint towards different pathways for *scyllo*-inositol utilization. It has been shown that *scyllo*-inositol levels can be altered independent from *myo*-inositol, e.g. in brain-derived glioma tumor models.^85^ We now provide [^13^C_6_]*scyllo*-inositol as a new tool that could be combined with other isotopic tracers for *myo*-inositol and glucose to investigate such distinct metabolic roles for inositol isomers. Together with the emerging field of lipidomics, these tracers provide the option for a more comprehensive understanding of inositol and PI(P) metabolism and signaling.

## Limitations of this study

One limitation of this study is the separation of *myo*- and *scyllo*-inositol by HILIC. The method provides sufficient separation of both isomers when used in similar concentrations. In the experiments, we often had a high concentration of one isomer and a low concentration of the other. Tailing or fronting of the much more concentrated peak (> 150x) into the smaller peak limited the accuracy for detection and quantification.

The NMR and lipidomics studies for identifying [^13^C_6_]PI were limited by there being no *scyllo*-PI standard available. Therefore, we were not able to unambiguously assign the NMR signals to [^13^C_6_]*scyllo*-PI species.

While the experiments provide evidence for the existence of [^13^C_6_]*scyllo*-PI species in mammalian cells, follow-up experiments are needed for a better understanding of their biological role.

## Acknowledgement

We acknowledge Dave Palmer for his input on the synthesis strategy and Hanaa Al Beiruty for synthetic assistance. We are also grateful to the entire Fiedler lab for proofreading the manuscript. Meike Marie Amma and Dorothea Fiedler acknowledge funding by the DFG grant 278001972-TRR186 TP24. Laxmikanth Kollipara and Sven Heiles acknowledge the support by the “Ministerium für Kultur und Wissenschaft des Landes Nordrhein-Westfalen”, „Senatsverwaltung für Wissenschaft, Gesundheit und Pflege Berlin“ and by the „Bundesministerium für Forschung, Technologie und Raumfahrt”. Adolfo Saiardi acknowledges the support of the Medical Research Council grant MR/T028904/1.

## Experimental section

### General Information

Chemicals were obtained from Sigma Aldrich, VWR, Roth, TCI, Thermo Scientific, or Roche and used without further purification unless stated otherwise. [^13^C_6_]D-glucose with an isotopic purity of 99% and all deuterated solvents were purchased from Eurisotope.

NMR spectra of synthesized molecules were recorded on a Bruker AV-III spectrometer operating at 600 MHz for proton nuclei or 151 MHz for carbon nuclei using a 5 mm RT-BBFO probe equipped with one-axis self-shielded gradients. NMR data are given as follows: chemical shift δ in ppm (multiplicity, coupling constant(s) J Hz, relative integral), where multiplicity is defined as: s = singlet, d = doublet, t = triplet, q = quartet, m = multiplet, br = broad or combinations of the above.

The software used to control the spectrometers was TopSpin 3.5 pl6.

For lipidomics experiments following chemicals were purchased from Sigma-Aldrich, Germany: ammonium formate, EquiSPLASH™ deuterated standards (Avanti), methyl tert-butyl ether, and formic acid. Precellys homogenization kit (2 mL prefilled with CK14 beads) and Precellys Evolution Touch + Cryolis Evolution Combo tissue homogenizer from BioCat GmbH, Germany. All ULC solvents, such as acetonitrile, isopropanol and methanol were purchased from Biosolve, Valkenswaard, the Netherlands.

## Mammalian cell culture and isotopic tracing

### Cultivation of cells

A172 and HCT116 cells were purchased from ATCC. HEK293T cells were obtained from Leibniz Institute DSMZ-German Collection of Microorganisms and Cell Cultures GmbH. HEK293T and HCT116 cells were cultivated in high glucose DMEM with GlutaMAX™ (Gibco™ #61965026). A172 cells were cultivated in high glucose DMEM, with GlutaMAX™ and pyruvate (Gibco™ #31966021). All cell media were supplemented with streptomycin/penicillin (final concentration of 100 U/mL, Gibco™ #15140122) and 10% FBS (Gibco™ #A5256801). Cells were cultivated at 37 °C in an atmosphere containing 5% CO_2_ and 95% humidity. PBS (#14190094) and trypsin (#12605010) were purchased from Gibco™, ThermoFisher Scientific.

### Media for isotopic tracing

Custom high glucose DMEM (Gibco™ #ME17481P1) with no regular inositol and 10% dialyzed FBS (Gibco™ #26400044), was used for all labeling experiments. For A172 cells, 1 mM sodium pyruvate was used as an additive. The synthesized inositol isomers were dissolved in ddH_2_O, sterile filtered, and stored as 50 mM stock solutions.

### Cell proliferation

Shortly, cells were seeded, and the medium was exchanged after 24 h with media containing different inositol concentrations and isomers. The following cell numbers were seeded: A172 (1,000 cells/well), HCT116 (400 cells/well) and HEK293T (1,500 cells/well). For the cell proliferation assay, Cell Proliferation Reagent WST-1 from Roche was used. The assay was performed as described in the protocol of the manufacturer.

### Isotopic tracing of [^13^C_6_]*scyllo*-inositol – A172 uptake experiments

A172 cells were seeded in 6-well plates in 3 mL medium for isotopic tracing with either 100 µM [^13^C_6_]*myo*-inositol, 100 µM [^13^C_6_]*scyllo*-inositol or a combination of 90 µM [^12^C_6_]*myo*-inositol and 10 µM [^13^C_6_]*scyllo*-inositol. For t=0, 24 and 48 h, 50,000 cells/well were seeded. For t=72 and 96 h, 25,000 cells/well were seeded. Additionally, a 15 cm dish with 3×10^5^ cells in the respective labeling media was seeded. After 5 days, the dishes were used to seed into the 6-well plates for the next five days, and also for another 15 cm dish, which was then used on day 10 for seeding the cells for the next five days. Before harvesting, cells were washed twice with 3 mL PBS and then harvested by trypsination (0.4 mL, for 5 min at 37 °C). The trypsin was quenched with 0.8 mL ice-cold PBS, and the cell suspension was transferred into 1.5 mL test tubes. After centrifugation for 5 min at 18,000 *g* and 4 °C, the supernatant was aspirated, and the cell pellets were flash-frozen and stored at −80 °C until further use.

### Sample preparation for the measurement of intracellular inositols

The cell pellet was thawed on ice, resuspended in MeOH/MeCN/ddH_2_O (40:40:20, v/v/v) by vortexing for 0.5 min and then incubated in a thermoshaker for 10 minutes at 80 °C, 500 rpm. After centrifugation (18,000 *g*, 10 min, 4 °C), the supernatant was quantitatively transferred into a glass-vial for HILIC-MS/MS analysis, and the pellets were used for the determination of the protein concentration *via* BCA. Exact volumes for the respective experiments are listed in Table S4.

### Determination of protein concentration by BCA

The cell pellets were resuspended in 1% SDS in PBS by pipetting. The suspensions were incubated for 20 min at 37 °C and 500 rpm. For the determination of the protein concentration, a Pierce™ BCA Protein Assay Kit (Thermo Scientific™) was used, and the assay was performed as described in the protocol of the manufacturer. Volumes used for resolubilization for the respective experiments are listed in Table S4.

### HILIC-MS/MS analysis of intracellular inositol

Analysis was performed on a Q Exactive Orbitrap Mass Spectrometer (Thermo Fisher Scientific Inc., Bremen, Germany) with a heated electrospray ionization (HESI) source. The Q Exactive was coupled to an UltiMate 3000 UHPLC system (Thermo Fisher Scientific Inc., Germering, Germany). For chromatographic separation, a method described in the literature was adapted.^1^

Mobile phase consisted of 50% MeCN with 0.04% NH_4_OH (buffer A) and 90% MeCN with 0.04% NH_4_OH (buffer B). The injection volume was 1 µL.

The elution started with a flow of 0.4 mL/min with 91% B held for 1.5 min and then decreased to 82.2% B between 1.5 min and 22.0 min. For rinsing the column, the flow was then reduced to 0.25 mL/min, and B was further decreased to 50% between 22.0 min and 25.0 min. Between 25.0 min and 30.0 min, B was held at 50%, but the flow was further reduced to 0.2 mL/min. For re-equilibration of the column, B was increased to 91% between 30.0 min and 33.0 min at a flow of 0.2 mL/min, then only the flow was also increased again to 0.35 mL/min between 33.0 and 35.0 min and even further to 0.4 mL/min between 35.0 min and 40.0 min. These conditions were held for another 5.0 min. The sample vials were maintained at 4 °C in an autosampler. Injection volume was 1 µL.

MS detection was performed using parallel reaction monitoring (PRM) and negative electrospray ionization. The run time was between 3.0 min and 22.0 min of the HILIC-MS/MS method. The HESI source parameters were set as follows: spray voltage was set at 2.5 kV; sheath gas flow rate at 60 (arbitrary units); Aux gas flow rate at 30 (arbitrary units), Sweep gas flow rate at 0 (arbitrary units); capillary temperature at 300 °C, S-lens RF level at 55.0 and the Aux gas heater temperature at 300 °C. The HESI probe depth was set to ring position C. The MS parameters were set as follows: the resolution was set to 17,500; the AGC target was set to 2e^5^, the maximum IT was set to 120 ms, and the isolation window was set to 1.0 *m/z*. Normalized collision-energy was set to 45. The inclusion list consisted of two entries for [^12^C_6_]inositol isomers 179.0560 *m/z* and for [^13^C_6_]inositol isomers 185.0760 *m/z*.

For quantification, external calibration curves with [^13^C_6_]*scyllo*-inositol and [^12^C_6_]*myo*-inositol were measured on the same day. [^13^C_6_]*scyllo*-inositol was also used for the quantification of [^12^C_6_]*scyllo*-inositol, and [^12^C_6_]*myo*-inositol was also used for the quantification of [^13^C_6_]*myo*-inositol. At least ten different concentrations were used for the calibration curve between 400 µM and 25 nM. For the 15-day uptake experiment, the measured concentrations were 400 µM, 200 µM, 100 µM, 50 µM, 10 µM, 5 µM, 1 µM, 500 nM, 250 nM, 100 nM, 75 nM, 50 nM, 25 nM. For intracellular inositols and hydrolyzed headgroups after treatment of A172, HEK293T and HCT116, the measured concentrations were 400 µM, 200 µM, 100 µM, 50 µM, 25 µM, 12.5 µM, 6.25 µM, 3.13 µM, 1.56 µM, 781 nM, 391 nM, 195 nM, 97.7 nM, 48.8 nM and 24.41 nM.

A weighted (1/x^2^) linear regression was used as a calibration model. For quantification of [^13^C_6_]isomers, the fragment with *m*/*z* 90.0180 and [^12^C_6_]isomers, the fragment with *m*/*z* 87.0077 were used, both with a mass tolerance of ± 5 ppm. For data analysis, FreeStyle™ 1.8 SP2, Version 1.8.63.0 (Thermo Fisher Scientific Inc.) and GraphPad Prism Version 10.5.0 were used. MS^2^ spectra of ^12^C and ^13^C_6_ *myo*-Inositol, Chromatographic separation and external calibration curves are shown in Figure S2. Robustness and signal-to-noise of the lower limit of quantification (LOQ) are listed in Table S3.

### Isotopic tracing of [^13^C_6_]*scyllo*-inositol – Lipid incorporation and intracellular inositol A172, HEK293T and HCT116

Cells were seeded at a cell type-dependent density in a 15 cm dish and cultured for two passages in media for isotopic tracing with 100 µM [^13^C_6_]*scyllo*-inositol. A172 cells were seeded at a density of 6×10^5^ in a 15 cm dish. HCT116 and HEK293T cells were seeded at a density of 3×10^5^ in a 15 cm dish. Cells were split at a confluency of around 85%. Once reaching again around 85% confluency, the cells were washed twice with PBS, harvested by trypsination (3 mL, for 5 min at 37 °C). The trypsin was quenched with 10 mL ice-cold PBS, and the cell suspension was transferred into 15 mL tubes. After centrifugation for 5 min at 200 g, the cell pellet was resuspended in 1.5 mL ice-cold PBS and transferred into precooled 2 mL test tubes. After centrifugation for 5 min at 200 *g* at 4 °C, the supernatant was aspirated, and the cell pellets were flash-frozen and stored at −80 °C until further use.

### Lipid extraction followed by hydrolysis for inositol isomer phosphatidylinositol headgroup measurement

The lipid extraction protocol was adapted from a literature-known procedure and optimized to investigate the inositol isomer headgroup composition of cells treated with 100 µM [^13^C_6_]*scyllo*-inositol.^63^ The cell pellet was thawed on ice and resuspended in 66 µL ddH_2_O. 250 µL MeOH was added, and the samples were first vortexed for 30 s and then further shaken for 20 min at 1000 rpm and at 4 °C. Then, 500 µL CHCl_3_ was added, vortexed for 30 s, and then shaken for 5 min at 1000 rpm and at 4 °C. Afterwards, 125 µL 6% metaphosphoric acid and 25 µL KClO_4_ solution (saturated, pH 6.5) were added and vortexed for 30 s. The sample was then centrifuged for 15 min at 18,000 *g* and 4 °C. Centrifugation resulted in phase separation with the CHCl_3_ phase on the bottom, aq. MeOH phase on top, protein as a thin layer in between. Only the CHCl_3_ phase was further processed. The CHCl_3_ layer was transferred to a 50 mL centrifuge tube, and 1 mL ddH_2_O and 25 µL KClO_4_ (saturated pH 6.5) were added, vortexed for 30 s, and centrifuged for 5 min at 5,000 *g* and 4 °C. The aq. phase was transferred to a fresh 50 mL centrifuge tube. For the next aqueous washing steps, 25 µL KClO_4_ (saturated, pH 6.5) was always added. The aqueous wash of the CHCl_3_ layer was repeated first with 2 mL ddH_2_O, then twice with 3 mL ddH_2_O, and in the end with 4 mL ddH_2_O, but in the last step, 250 µL CHCl_3_ was also added. The CHCl_3_ layer was transferred into a fresh 1.5 mL test tube and concentrated at 55 °C in a heat block (550 rpm). After evaporation, a visible residue remained at the bottom. For lipid hydrolysis, the residue was resuspended in 200 µL 9.2 M HClO_4_. The resuspension was incubated for 16 h at 80 °C and 1000 rpm. The sample was then put on ice and neutralized with 10 M KOH. The pH was adjusted to 7.0 ±0.5 with a final volume of 1 mL. The suspension was centrifuged for 10 min at 18,000 *g* and 4 °C, and the supernatant was transferred into a new 2 mL test tube. The solvents were removed by lyophilization, and the sample was stored at −20 °C until HILIC-MS/MS measurement. For measurement, the lyophilized supernatant was redissolved in 100 µL MeOH/MeCN/ddH_2_O (40:40:20, v/v/v), centrifuged for 10 min at 18,000 *g* and 4 °C, and the resulting supernatant was transferred into a glass vial for HILIC-MS/MS analysis. For protein normalization, sister plates were used.

### Lipidomics LC-MS/MS sample preparation

All samples, i.e. A172 (90 µM *myo*-inositol and 10 µM [^13^C_6_]*scyllo*-inositol), A172 (100 µM [^13^C_6_]*scyllo*-inositol), HEK293T (100 µM [^13^C_6_]*scyllo*-inositol) and HCT116 (100 µM [^13^C_6_]*scyllo*-inositol) were processed based on the liquid-liquid extraction (LLE) protocol by Matyash *et al*.^65^ Briefly, to each cell pellet, 225 µL of cold MeOH was added and after a brief vortex the suspension was transferred into a 2 mL prefilled tubes containing CK14 beads (1.4 mm zirconia beads). Next, 10 µL of undiluted internal standards mixture (EquiSPLASH) was added. In parallel, two extraction blanks, i.e., with and without 10 µL EquiSPLASH, were prepared in 2 mL prefilled tubes and processed in the same manner as below. The sample tubes were subjected to homogenization using Precellys Evolution Touch + Cryolis Evolution Combo at 4 °C, 6000 rpm, 4 × 15 s, 10 s pause. Next, 750 µL of cold MTBE was added, briefly vortexed, and homogenized as above. Next, samples were placed on vortex at 4 °C for 1 h (800 rpm). Phase separation was induced by adding 188 µL of ddH_2_O and centrifugation for 15 min, 16,000 *g* at 4 °C. The upper layer was carefully collected in a new LoBind Eppendorf tube (Extract 1). The middle layer was re-extracted with 500 µL of MTBE/MeOH/ddH_2_O (10:3:2.5, v/v/v), vortexed at 4 °C for 30 min (800 rpm), and centrifuged as above. Lastly, the upper layer was collected and combined with Extract 1 and dried under vacuum (SpeedVac). Dried lipid extracts were reconstituted in 100 µL of 100% isopropanol. For proper solubilization, sample tubes were placed in an ultrasonic water bath followed by centrifugation at 14,000 *g* (each for 3 min), and the supernatant was transferred into HPLC glass inserts before lipidomics LC-MS/MS analysis.

### Lipidomics LC-MS/MS analysis

Each sample (5 µL) was analyzed using a Vanquish Flex UHPLC system coupled to Exploris-240 MS (both Thermo Scientific) *via* HESI source. The separation was carried out on a C18 reversed phase 150 x 2.1 mm I.D., S-1.9 µm,12 nm column (YMC-Triart) using a binary gradient, i.e., buffer A: 85/10/5 IPA/MeCN/ddH_2_O (v/v/v) and buffer B: 50/50 MeCN/ddH_2_O (v/v), both containing 5 mM ammonium formate and 0.1% FA at final concentrations. The flow rate was 300 µL/min, and the column oven temperature was set to 50 °C. The LC gradient was as follows: 0-3 min (B 10%–25%), 3–9 min (B 25%–54%), 9–14 min (B 54%–77%), 14–18 min (B 77%–88%), 18–22 min (B 88%–95%), 22–30 min (B 95%), 30–30.5 min (B 95%–10%), 30.5–38 min (B 10%). The MS data were acquired in two different ways, i.e., data-dependent aquesition (DDA) and targeted-DDA (t-DDA). For both analyses, the ion source settings: positive ion mode and negative ion mode voltages were set to 3500 and 2500, respectively. Sheath, auxiliary, and sweep gas values were set to 40, 10, 1, respectively. RF% value was 40, ion transfer tube (ITT) and vaporizer temperature (VT) were kept at 250 °C and 300 °C, respectively, whereas for t-DDA, RF% was 50, ITT 300 °C, and VT 350 °C were used. For DDA mode, MS survey scans were acquired in the Orbitrap from *m/z* 150 to 1500 at a resolution of 120000. Precursors were isolated with the Quadrupole at a 2 *m/z* window, and MS/MS analysis was performed using the TOPN approach (N=10) using higher energy collisional dissociation fragmentation with normalized collision energies at 25, 30, 35%. MS/MS spectra were acquired at a resolution of 15,000. A dynamic exclusion of 10 s with low/high mass tolerance of 5 ppm was applied. Automatic gain control (AGC) target and intensity threshold values were set to 1 × 10^6^ and 5 x 10^3^ for MS and AGC value 1 × 10^5^ for MS/MS scans. Maximum injection times were set to auto for both, and all data was recorded in profile mode.

The t-DDA MS analysis was based on the database search results of the untargeted DDA runs that were acquired in the negative ion mode. MS raw data was first processed using the Lipostar2 software.^66^ The precursor *m/z* values of all [^12^C_6_]PI species were extracted, manually added 6.02013 Da, and created a putative list of [^13^C_6_]PI species. This was used as a “targeted list” with and without [^12^C_6_]PI masses and remeasured all samples using the following settings. For DDA mode, MS survey scans were acquired in the Orbitrap from *m*/*z* 150 to 1500 at a resolution of 120000. Precursors were isolated with the Quadrupole at 1.6 *m*/*z* window, and MS/MS analysis was performed using TOP speed approach (cycle time 3 s) using higher energy collisional dissociation fragmentation with normalized collision energies at 20, 25, 30, 35, 45%. MS/MS spectra were acquired at a resolution of 15,000. Low/high mass tolerance set to ± 10 ppm. AGC target value was set to standard for MS and 1 × 10^5^ for MS/MS scans. Maximum injection times were set to auto for MS and dynamic for MS/MS, respectively. The time mode was set to unscheduled.

### Lipidomics data analysis using Lipostar2 software

The DDA MS raw data were imported into Lipostar2.1.4, and the vendor-provided “Thermo DDA Negative” default settings were applied, including instrument and peak picking. Next, data filtering criteria were: mass tolerance of ± 5 ppm; retention time tolerance of 0.20 min; and only precursors with isotopic peak pattern and MS/MS spectra were allowed for identification. The filtered masses were searched using the Lipid Maps database downloaded on 20.02.2025. After the identification process, only those lipid species that qualified “automatic approval” (4 and 3 stars) were exported and used for further analysis, such as manually validating the [^12^C_6_]PI/[^13^C_6_]PI MS and MS/MS signals in the raw data and generating the putative list of [^13^C_6_]PI species for t-DDA analysis.

### Lipidomics determination of relative abundance and identification of [^13^C_6_]PI

For further data analysis, FreeStyle™ 1.8 SP2, Version 1.8.63.0 (Thermo Fisher Scientific Inc.) was used. For the relative abundance of PIs, the data of the t-DDA with the [^12^C_6_]PI/[^13^C_6_]PI method was used, if at least five MS^1^ spectra were detected in at least two of the three replicates. The peak height of the described [^12^C_6_]/[^13^C_6_]PIs was plotted. For correction of the sample preparation, the deuterated PI contained in the added EquiSPLASH was used as an internal standard. The samples were normalized by the protein concentration *via* BCA. The [^12^C_6_]PIs were assigned with LipoStar2. A signal was assigned as putative [^13^C_6_]PI if it was co-eluting with its Lipostar2 assigned [^12^C_6_]PI, had an added mass shift of 6.02013 Da, and had at least five MS^1^ spectra in at least two of the three replicates.

For identification of [^13^C_6_]PIs, at least one MS^2^ spectrum in at least two of the three replicates of the samples measured with the t-DDA [^13^C_6_]PI only method had to be detected. First, all MS^2^ spectra that contained the fragment of the [^13^C_6_]PI headgroup (*m/z* 247.0320 Da) were assigned to the putative PI based on retention time and precursor ion. For validation of the identification, a transition list was created in Skyline with the tool LipidCreator for all putative [^13^C_6_]PI with a MS^2^ spectrum that contained the [^13^C_6_]PI headgroup (*m/z* 247.0320 Da) fragment. For the assignment “identified”, the mentioned MS^2^ spectra had to contain, next to the precursor ion, the fragments of glycerophosphate (*m/z* 152.9958 Da), [^13^C_6_]PI headgroup (*m/z* 247.0320 Da), and the fragments of the different fatty acyls, all with a mass tolerance of ± 5 ppm.

### NMR lipid extraction

The whole lipidomics workflow described above, starting from metabolic labeling, was carried out for A172 cells cultivated in either 100 µM [^13^C_6_]*scyllo*-inositol, [^13^C_6_]*myo*-inositol, [^12^C_6_]*myo*-or [^12^C_6_]*scyllo*-inositol. Five cell pellets were combined per measurement. The lipid extraction was adapted from the workflow described in the lipidomic-LC-MS/MS workflow. Instead of a Precellys Evolution Touch + Cryolis Evolution Combo, samples were placed in an ultrasonic water bath filled with ice water for homogenization. The organic layer was concentrated under N_2_ flow and transferred into a fresh 1.5 mL test tube and concentrated at 55 °C in a heat block (550 rpm). The residue was redissolved in 480 µL CDCl_3_ and 120 µL CD_3_OD (4:1), centrifuged for 10 min at 18,000 g and 4 °C, and the resulting supernatant was transferred into a NMR tube. The sample was stored at −20 °C until measurement.

### NMR measurement

Measurements of the metabolic extracts were performed at 300 K on a Bruker AV-III spectrometers (Bruker Biospin, Rheinstetten, Germany) operating at 600 MHz for proton nuclei or 151 MHz for carbon nuclei using a cryogenically cooled 5 mm QCI-triple resonance probe equipped with one-axis self-shielded gradients. For measurement, processing, and analysis of the NMR datasets, TopSpin 3.5 pl6 was used. For calculating the P1 pulse, the samples were manually locked, matched, and shimmed for each sample. To record [^1^H, ^13^C]-HSQC and [^1^H, ^13^C]-HSQC-CLIP-COSY spectra, home-written pulse programs were used. The experiements utilized a BIRD-Puls to suppress signals of protons not bound to ^13^C.^67,86^ 512 complex points were recorded, and the number of scans was kept at 768. The spectral width was set to 10,000 Hz (center at 4.7 ppm) and 9,090 Hz (center at 70 ppm) for ^1^H and ^13^C, respsctively. All the spectra were processed without digital water suppression with manual phase correction and automatic baseline correction.

## Synthesis

### [^13^C_6_]*myo*-inositol (1R,2S,3R,4S,5R,6S)-Cyclohexan-1,2,3,4,5,6-hexol)

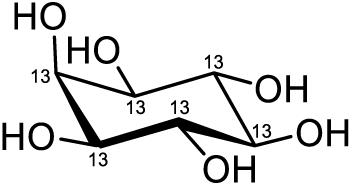

[^13^C_6_]*myo*-inositol was synthesized according to (*Chem. Sci*., **2019**, ***10***, 5267-5274), whereby the comments published in (*ACS Cent. Sci*. **2022**, *8*, 1683−1694) were considered.^40,42^ Additionally, the following changes were made:

Toluene was used for the co-evaporation of pyridine to further minimize the amount of pyridine during the workup of [^13^C_6_]*myo*-inositol hexakisacetate. The concentrate was resuspended in CH_2_Cl_2_ and 1 M HCl. In the next step, the suspension was filtered through a cotton plug instead of a paper filter. Following the extraction, the crude product was purified *via* flash column chromatography. A known yellow impurity was monitored closely, and only slightly yellow fractions containing [^13^C_6_]*myo*-inositol were used for the next step.

### Glycerol stock of *Gluconobacter oxydans* 2343

To prepare the growth medium, 25.0 g D-sorbitol and 1.25 g yeast extract were dissolved in 250 mL ddH_2_O and sterile filtered. For the preparation of glycerol stocks, all steps were done next to the flame. *Gluconobacter oxydans* was purchased from Leibniz Institute DSMZ-German Collection of Microorganisms and Cell Cultures GmbH as a lyophilized powder and resuspended in 0.5 mL growth medium. The suspension was mixed gently by pipetting, transferred to a test tube containing 6 mL of growth medium, and incubated on an orbital shaker at 30 °C, 220 rpm for 21 h. The OD_600_ of the primary culture was 2.8. For the glycerol stocks, a one-to-one mixture of the bacterial culture and 50% autoclaved glycerol in ddH_2_O was prepared, aliquoted in 1 mL, and frozen and stored at −80 °C.

### [^13^C_6_]*scyllo*-inosose ((2R,3S,4S,5R,6S)-2,3,4,5,6-pentahydroxycyclohexan-1-one)

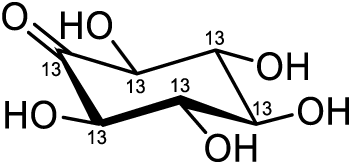

The synthesis was adapted from literature reports.^49^ All steps until the centrifugation were done next to the flame. The same growth medium was used as described for the preparation of the glycerol stock. For the labeled synthesis medium, [^13^C_6_]*myo*-inositol (600 mg, 3.22 mmol), D-sorbitol (20.0 mg), and yeast extract (100 mg) were dissolved in 20 mL ddH_2_O and filtered under sterile conditions. For reaction control, a synthesis medium with non-labeled *myo*-inositol was prepared. For the growth culture, two scratches of the glycerol stock were added to 50 mL growth medium in a 250 mL autoclaved Erlenmeyer flask and incubated on an orbital shaker at 30 °C, 220 rpm for 20 h. 2.5 mL of growth medium was added to 20 mL of the labeled synthesis medium in a 100 mL autoclaved Erlenmeyer flask. For NMR reaction control, 1 mL of the control culture was spun down (20 min, room temperature, 18000 *g*), and the supernatant was filtered. 500 µL of the filtrate was combined with 60 µL D_2_O, and a ^1^H NMR with 512 scans was measured. For a LC-MS reaction control, 5 µL of the control culture was added to 495 µL ddH_2_O and filtered. 50 µL of the filtrate was further diluted with 150 µL 80% MeCN in ddH_2_O and measured *via* LC-MS in HILIC mode. Both synthesis cultures were incubated for 72 h until no *myo*-inositol could be detected by NMR. If no *myo*-inositol was detected anymore, the culture was further incubated for 4 h. Cells were removed by centrifugation (18,000 *g*, 30 min, 10 °C). The cell pellet was washed once with 10 mL dH_2_O. The supernatant was concentrated to 5 mL under reduced pressure. The concentrate was transferred into a 50 mL centrifuge tube, and the product was precipitated with 45 mL MeCN overnight at 4 °C. The centrifuge tube was centrifuged (4 °C, 3214 *g,* 30 min), the supernatant was decanted and the precipitate was washed three times with 5 mL ice-cold dH_2_O. The supernatant was combined with the supernatant from the wash steps, concentrated to 5 mL under reduced pressure, and another round of precipitation was carried out as described before. The combined precipitates were dried *via* lyophilization to yield [^13^C_6_]*scyllo*-inosose (540 mg, 2.93 mmol, 91%) as a white powder. It is described in the literature that inosose is in a complex equilibrium between its keto, enol, and hydrated forms. Therefore, the NMR spectra contained more than signals than expceted.^87,88^

**^1^H-NMR (decoupled):** (600 MHz, D_2_O) *δ* = 4.51 – 4.41 (m, 2H), 3.92 – 3.81 (m, 1H), 3.49 (qd, *J* = 9.0, 8.2, 4.9 H, 5H), 3.42. (t, J = 8.5 Hz, 1H).

**^13^C-NMR:** (150 MHz, D_2_O) δ = 206.83 – 205.35 (m), 94.96 – 93.37 (m), 76.89 – 75.14 (m), 74.77 – 71.54 (m).

**HRMS (ESI):** calc. for ^13^C_6_H_10_O_6_ [M−H]^−^: 183.0606, found: 183:0605.

### [^13^C_6_]*scyllo*-inositol ((1R,2R,3R,4R,5R,6R)-cyclohexane-1,2,3,4,5,6-hexol)

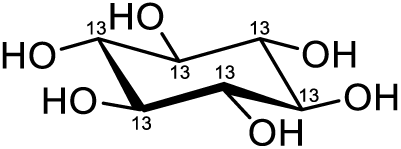

[^13^C_6_]*scyllo*-inosose (100 mg, 0.543 mmol, 1.0 equiv.) was dissolved in 20 mL dH_2_O by heating the mixture to 80 °C for 30 min. After *scyllo*-inosose was completely dissolved, the solution was cooled down to room temperature, and 180 mL MeCN was added. After 10 min, sodium triacetoxyborohydride (345 mg, 1.629 mmol, 3.0 equiv.) was added to a vigorously stirred clear solution. Formation of precipitate could be observed upon the addition, and the suspension was stirred at room temperature for 1 h. For reaction control, a 50 µL sample was diluted with 200 µL 50% MeCN in ddH_2_O with 0.1% NH_3_ and measured *via* LC-MS in HILIC mode. After full conversion, the suspension was concentrated under reduced pressure. The crude product was dissolved in 50 mL dH_2_O by heating the mixture to 70 °C for 20 min and filtered through a plug of DOWEX^®^ 50W x 8 (H^+^ form; 3 cm height, 7 cm diameter). The filter bed was further washed with 200 mL of sequentially added dH_2_O. The filtrate was concentrated to 5 mL under reduced pressure and precipitated with 45 mL MeCN. The precipitate was dried by lyophilization to obtain a mixture of [^13^C_6_]*scyllo*-inositol and [^13^C_6_]*myo*-inositol. The separation of the isomers was performed with an adapted procedure from the literature.^51^ The column volume of the anion exchange column (DOWEX^®^ 1 X 8) was 60 mL. The column was activated with 200 mL sodium tetraborate (100 mM; 40 mL 0.5 M boric acid mixed with 60 mL 1 M NaOH and 100 mL dH_2_O) and washed with 200 mL dH_2_O. The mixture of [^13^C_6_]*scyllo*-inositol and [^13^C_6_]*myo*-inositol was dissolved in 20 mL dH_2_O by heating the mixture to 70 °C for 30 min. The clear solution was cooled down to room temperature. Boric acid (600 µL, 0.5 M) was added to derivatize [^13^C_6_]*myo*-inositol, and the solution was stirred for 30 min before loading on the prepared anion exchange column. To obtain pure [^13^C_6_]*scyllo*-inositol, 250 mL dH_2_O was used as the mobile phase. For the first fraction, 50 mL were collected, then the fraction size was switched to 7 mL. It is important to note that the column must never be stopped during the elution of [^13^C_6_]*scyllo*-inositol; otherwise, co-elution of [^13^C_6_]*myo*-inositol was observed. The fractions were monitored *via* LC-MS/MS in HILIC mode. Fractions containing only [^13^C_6_]*scyllo*-inositol were combined. 100 µL formic acid was added, the combined fractions were concentrated to 30 mL, and filtered through a plug of Amberlite™ MB-6113 Ion Exchange Resin (height 2 cm, diameter 4 cm). The filter bed was further washed with 150 mL of sequentially added dH_2_O. The filtrate was concentrated to 5 mL under reduced pressure and precipitated with 45 mL MeCN. The precipitate was dried by lyophilization to yield [^13^C_6_]*scyllo*-inositol (35 mg, 0.188 mmol, 35%) as a white powder with an isotopic purity of 99%.

**^1^H-NMR (decoupled):** (600 MHz, D_2_O) δ = 3.38 (s, 6H)

**^13^C-NMR:** (150 MHz, D_2_O) δ = 73.4 (C 1-6)

**HRMS (ESI):** calc. for ^13^C_6_H_12_O_6_ [M−H]^−^: 185:0762, found: 185.0761

### HILIC-MS analysis for reaction control

Reaction controls were performed on an Agilent 1260 Infinity Binary LC coupled to an Agilent 6130 Single Quadrupole LC-MS System (Agilent, Waldbronn, Germany) with an electrospray ionization (ESI) source. The sample vials were maintained at room temperature in an autosampler. Hydrophilic interaction liquid chromatography was performed using a 2.1 x 50 mm, 2.6 μm Thermo Scientific™ Accucore™ 150 Amide-HILIC column. Column oven temperature was kept at 30 °C.

Used mobile phases consisted of 50% MeCN with 0.04% NH_4_OH (buffer A) and 90% MeCN with 0.04% NH_4_OH (buffer B). The injection volume was 1 µL.

The elution started with a flow of 0.7 mL/min with 100% B and decreased to 83% B in 1.0 min, between 1 min and 6.5 min, B was further decreased to 80%, and then reduced to 10% for 1 min. For equilibration of the column, B was again set to 100%.

MS detection was performed using negative electrospray ionization. The ESI source parameters were set as follows: spray voltage was set at 3 kV; drying gas flow at 12.0 L/min, nebulizer pressure at 55 psig; drying gas temperature at 300 °C. The peak width was set at 0.100 min, and the cycle time was 0.60 sec/cycle. The quadrupole was operated in selected ion monitoring (SIM) mode.

For [^12^C_6_]inositol, it was set as signal one with *m/z* 179.00 and *scyllo*-inosose as signal two with *m/z* 177.00. For [^13^C_6_]inositol, it was set as signal one with *m/z* 185.00 and *scyllo*-inosose as signal two with *m/z* 183.00. A gain of 2.00, a dwell time of 290 msec, and 50% cycle time were used.

Peak detection, integration, and quantification were performed using OpenLab ChemStation Rev. C.01.07 SR3 (Agilent, Waldbronn, Germany).

