## Supplementary Tables and Figures for "Isotopic tracing of [^13^C_6_]*scyllo*-inositol uncovers its incorporation into phosphatidylinositols in mammalian cells"

### Table S1: Conditions tested for reduction of *scyllo*-inosose

| Solvent | Reduction agent | t[°C] | Comment |
|---|---|---|---|
| $CH_3CN$ | $NaBH_4$ | 50 | Mixture *scyllo*- *myo*-inositol (1:1.5) |
| $CH_3CN$, $H_2O$ (3:1, v:v) | $NaBH_4$ | 50 | Mixture *scyllo*- *myo*-inositol (1:2)<br>Only here a clear solution |
| $CH_3CN$ | $NaBH_4$ | 25 | Mixture *scyllo*- *myo*-inositol (1:1.5) |
| $CH_3CN$ | $NaBH_4$ | 0 | Mixture *scyllo*- *myo*-inositol (1:1.5) |
| EtOH | $NaBH_4$ | 25 | No reaction |
| MeOH | $NaBH_4$ | 25 | Unidentifiable product |
| MeOH, $H_2O$ (19:1, v:v) | $NaBH_4$ | 25 | Dissolved before in hot water<br>Mixture *scyllo*- *myo*-inositol (1:3.8) |
| MeOH, $H_2O$ (9:1, v:v) | $NaBH_4$ | 25 | Dissolved before in hot water<br>Mixture *scyllo*- *myo*-inositol (1:3.4) |
| MeOH, $H_2O$ (9:1, v:v) | $NaBH_4$ | 40 | Dissolved before in hot water<br>Mixture *scyllo*- *myo*-inositol (1:3) |
| $CH_3CN$ | $NaBH(CH_3COO)_3$ | 25 | No reaction |
| **$CH_3CN$, $H_2O$ (9:1**, v:v**)** | **$NaBH(CH_3COO)_3$** | **25** | **Dissolved before in hot water**<br>**Mixture *scyllo*- *myo*-inositol (1:1.2)** |
| $CH_3CN$, $H_2O$, AcOH(9:1:0.02, v:v:v) | $NaBH(CH_3COO)_3$ | 25 | Dissolved before in hot water<br>Mixture *scyllo*- *myo*-inositol (1:1.8) |
| $H_2O$ | Raney®-Nickel | 93 | Mixture *scyllo*- *myo*-inositol (1:1.9) |

**TableS2: starting conditions in media and additives**; due to tailing of high [$^{12}C_6$]*myo*-inositol, [$^{12}C_6$]*scyllo*-inositol not always measurable

| | [$^{12}C_6$]*myo*-inositol [µM] | [$^{12}C_6$]*scyllo*-insoitol [µM] | [$^{13}C_6$]*scyllo*-inositol [µM] | [$^{13}C_6$]*myo*-inositol [µM] |
|---|---|---|---|---|
| **FBS** | 1066.48 | 32.70 | n.d. | n.d. |
| **dFBS** | 7.13 | 0.01 | n.d. | n.d. |
| **DMEM normal** | 95.15 | 3.36 | n.d. | n.d. |
| **100 µM [$^{13}C_6$]*myo*-inositol DMEM** | 0.73 | 0.00 | n.d. | 76.86 |
| **100 µM [$^{13}C_6$]*scyllo*-inositol DMEM** | 1.08 | 0.03 | 81.34 | n.d. |
| **90 µM [$^{12}C_6$]*myo*-inositol, 10 µM [$^{13}C_6$]*scyllo*-inositol DMEM** | 62.44 | 0.00 | 8.20 | n.d. |
| **DMEM w/o inositol** | 1.42 | 0.01 | n.d. | n.d. |

**Table S3: Basic information about the HILIC-MS/MS method**: interday robustness for 25 to 50,000 nM for 7 days sequentially, on the last day two measurements; for 100,000 to 400,000 nM for 5 days sequentially, signal to noise at 25 nM; no carry over in the method was observed

| **Variation coefficient** | Concentration [nM] | 25 | 50 | 100 | 500 | 1000 | 5000 | 10,000 | 50,000 | 100,000 | 200,000 | 400,000 |
|---|---|---|---|---|---|---|---|---|---|---|---|---|
| [$^{12}C_6$]*myo*-inositol *m/z* 87.0077 | | 14% | 13% | 10% | 10% | 10% | 7% | 5% | 7% | 5% | 6% | 5% |
| [$^{13}C_6$]*scyllo*-inositol *m/z* 90.0180 | | 15% | 15% | 11% | 10% | 11% | 9% | 6% | 7% | 6% | 6% | 6% |

| **Signal to noise at 25 nM** | | day 1 | day 2 | day 3 | day 4 | day 5 | day 6 | day 7-1 | day 7-2 |
|---|---|---|---|---|---|---|---|---|---|
| [$^{12}C_6$]*myo*-inositol *m/z* 87.0077 | | 16.8 | 35.4 | 36.2 | 96.6 | 53.8 | 29.6 | 84.1 | 84.1 |
| [$^{13}C_6$]*scyllo*-inositol *m/z* 90.0180 | | 23.8 | 85.2 | 45.9 | 57.4 | 20 | 85.1 | 16.4 | 39.2 |

### Table S4: Volumes for determination of intracellular inositol

| Experiment | Lysis volumes[µL] | Volumes for BCA [µL] | Comment |
|---|---|---|---|
| **15 d uptake in A172** | | | |
| t0 | 200 | 50 | |
| **100 µM [$^{13}C_6$]*myo*-inositol** | | | |
| 1 d | 100 | 50 | |
| 2 d | 200 | 50 | |
| 3 d | 100 | 75 | |
| 4 d | 200 | 75 | |
| 5 d | 200 | 75 | |
| 6 d | 100 | 50 | |
| 7 d | 200 | 50 | |
| 8 d | 100 | 75 | #8 lysis volume 200 µL |
| 9 d | 200 | 75 | |
| 10 d | 200 | 75 | |
| 11 d | 100 | 50 | |
| 12 d | 200 | 50 | |
| 13 d | 100 | 75 | |
| 14 d | 200 | 75 | |
| 15 d | 200 | 75 | |
| **90 µM [$^{12}C_6$]*myo*-inositol, 10 µM [$^{13}C_6$]*scyllo*-inositol** | | | |
| 1 d | 100 | 50 | |
| 2 d | 200 | 50 | |
| 3 d | 100 | 75 | |
| 4 d | 200 | 75 | |
| 5 d | 200 | 75 | |
| 6 d | 100 | 50 | |
| 7 d | 200 | 50 | |
| 8 d | 100 | 75 | |
| 9 d | 100 | 75 | |
| 10 d | 200 | 75 | |
| 11 d | 100 | 50 | |
| 12 d | 200 | 50 | |
| 13 d | 100 | 75 | |
| 14 d | 100 | 75 | |
| 15 d | 200 | 75 | |
| **100 µM [$^{13}C_6$]*scyllo*-inositol** | | | |
| 1 d | 100 | 50 | |
| 2 d | 200 | 50 | |
| 3 d | 100 | 75 | |
| 4 d | 200 | 75 | |
| 5 d | 300 | 75 | |
| 6 d | 100 | 50 | |
| 7 d | 200 | 50 | |
| 8 d | 100 | 75 | |
| 9 d | 200 | 75 | |
| 10 d | 300 | 75 | |
| 11 d | 100 | 50 | |
| 12 d | 200 | 50 | |
| 13 d | 100 | 75 | |
| 14 d | 100 | 75 | |
| 15 d | 100 | 75 | |
| **A172, HEK293T and HCT116 labeling for 8 d** | | | |
| A172 | 1000 | 750 | 1:2 dilution for [$^{13}C_6$]*scyllo*-inositol measurment |
| HEK293T | 500 | 750 | 1:2 dilution for [$^{13}C_6$]*scyllo*-inositol measurment |
| HCT116 | 500 | 750 | |

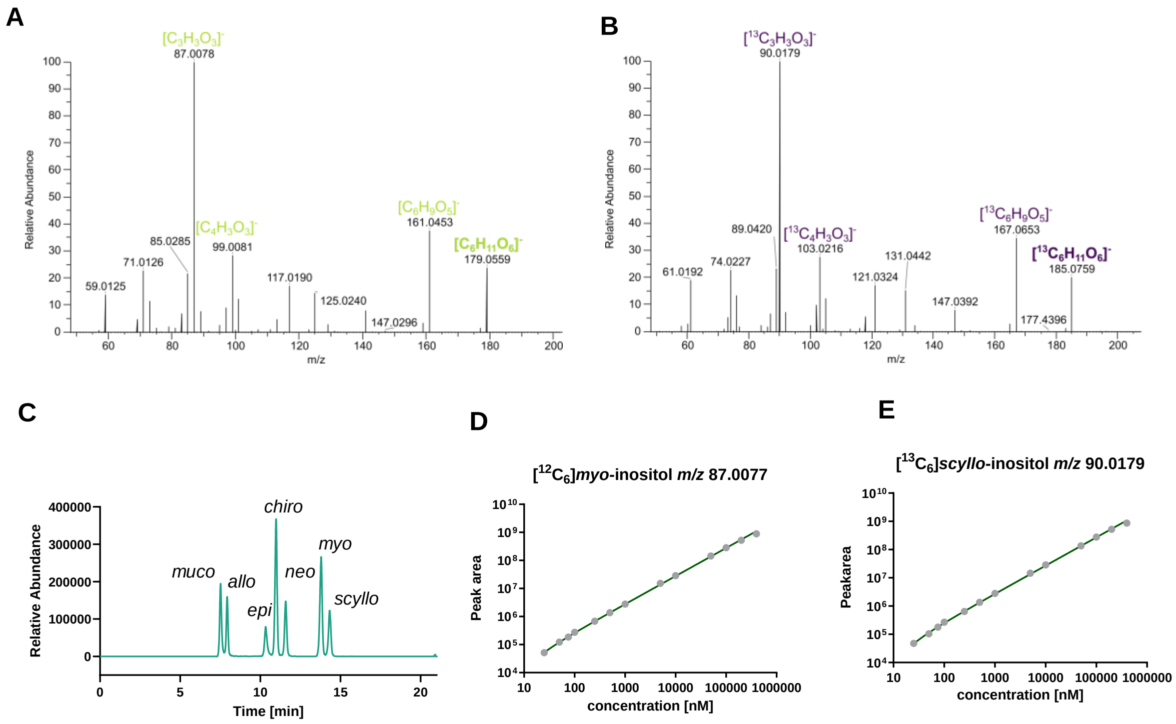


**Figure S1: HILIC-MS/MS methods**.

A–B) MS2 spectra for $[^{12}C_6]$*myo*-inositol (A) and $[^{13}C_6]$*myo*-inositol (B) with the respective precursor and three most abundant fragments. C) Chromatographic separation of seven inositol isomers. D–E) External calibration for $[^{12}C_6]$*myo*-inositol (D) and $[^{13}C_6]$sc*yllo*-inositol (E).

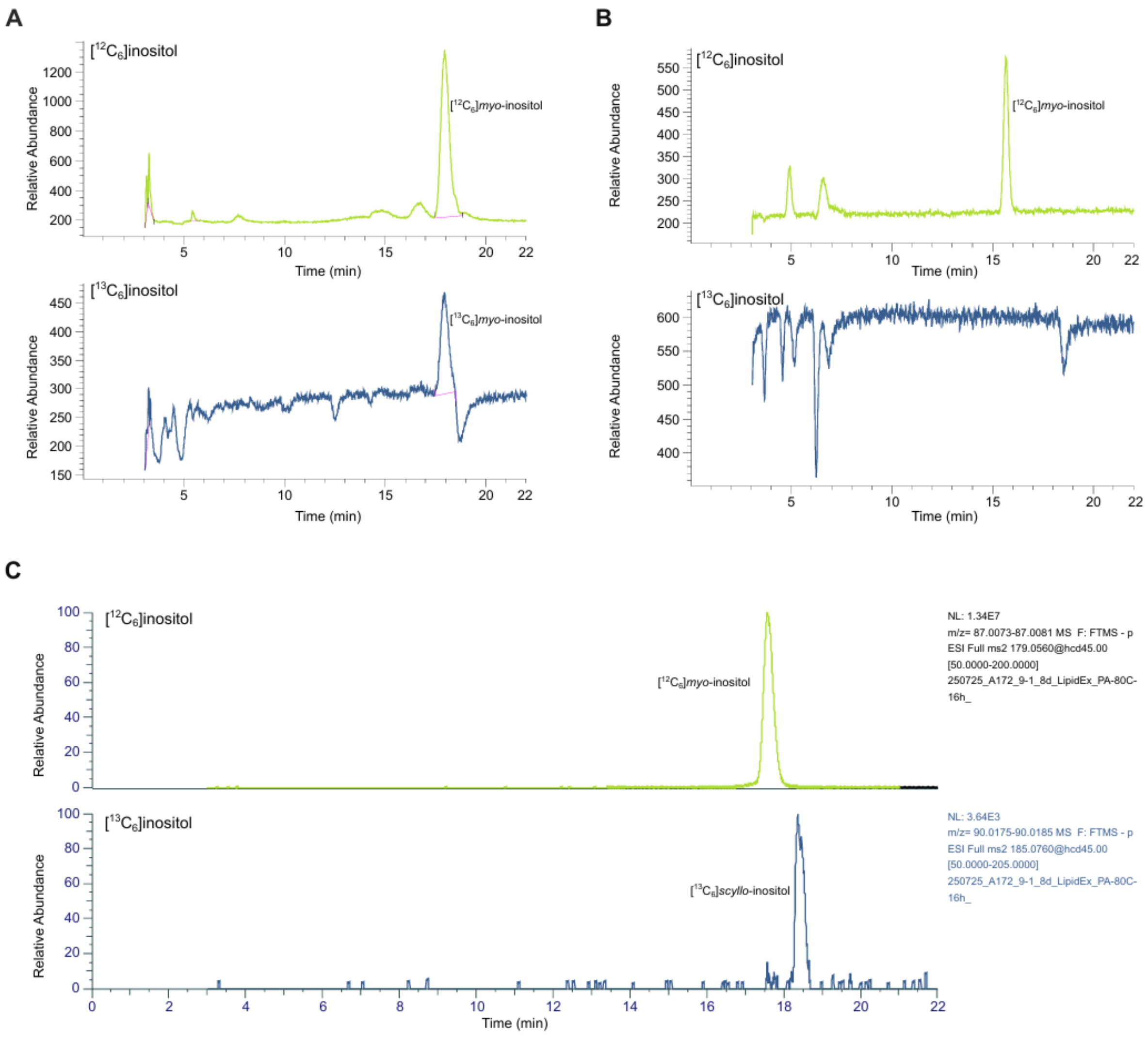


**Figure S2: Lipid extraction and hydrolysis of A172 cells.**

A–B) HILIC-MS/MS analysis of $[^{12}C_6]$ and $[^{13}C_6]$inositol isomers after lipid extraction and hydrolysis from A172 cells. The cells were cultured in normal DMEM and 100 nmol $[^{13}C_6]$*myo*-inositol were added prior to extraction to investigate carry-over of intracellular inositol. The initial conditions resulted in a significant carry-over (A), which could be prevented by an optimized washing procedure (B). C) Cells were cultivated in a combination of 90 µM $[^{12}C_6]$*myo*- and 10 µM $[^{13}C_6]$*scyllo*-inositol. 5x15 cm plates were combined for HILIC-MS/MS analysis to obtain a peak for $[^{13}C_6]$*scyllo*-inositol. Approx. 0.00007 nmol/mg total protein $[^{13}C_6]$*scyllo*-inositol was detected compared to 0.6154 nmol/mg total protein $[^{12}C_6]$*myo*-inositol.

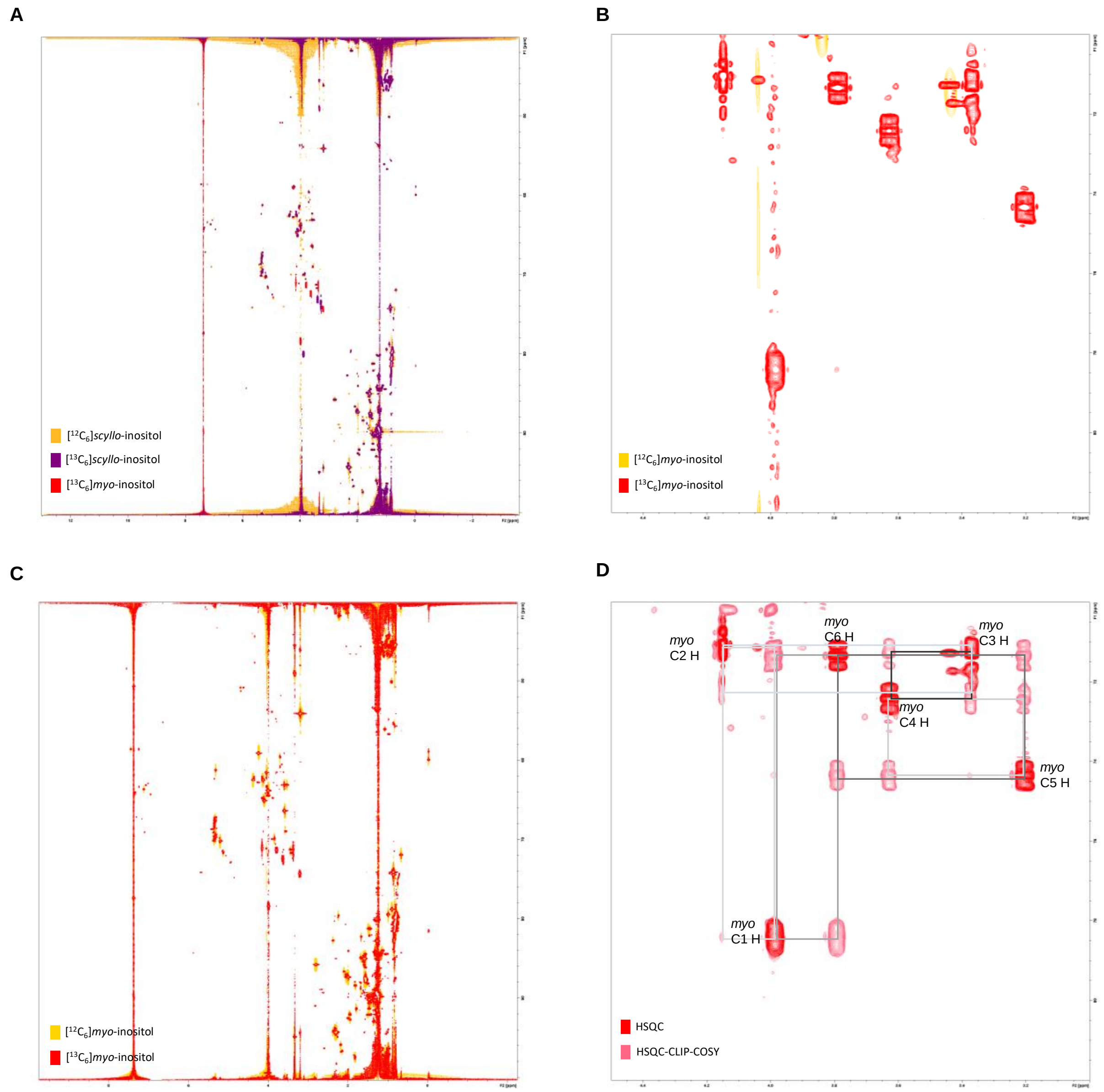


**Figure S3: 2D NMR spectra of lipid extracts from A172 cells**.
A) Zoomed out spectra shown in Figure 4B. B) [$^{1}$H,$^{13}$C]-HSQC-NMR of [$^{12}C_6$] and [$^{13}C_6$]*myo*-inositol. C) Zoomed out spectra shown in Figure S4B. D) [$^{1}$H,$^{13}$C]-HSQC-NMR and [$^{1}$H,$^{13}$C]-HSQC-CLIP-COSY of [$^{13}C_6$]*myo*-inositol.

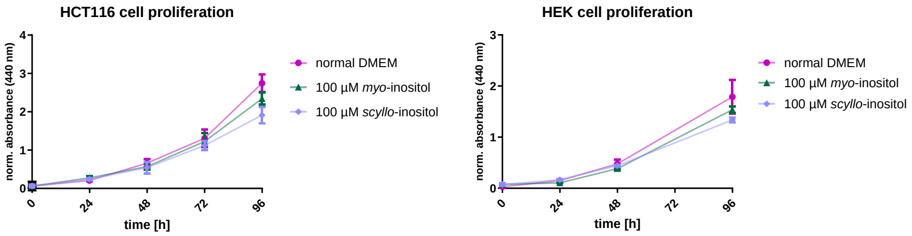


**Figure S4: Proliferation of HCT116 and HEK293T cells**

Cell proliferation of cells cultivated in defined inositol concentrations by a WST assay.

**NMR spectra of synthesized compounds**

**[$^{13}C_6$]*Scyllo*-inositol**

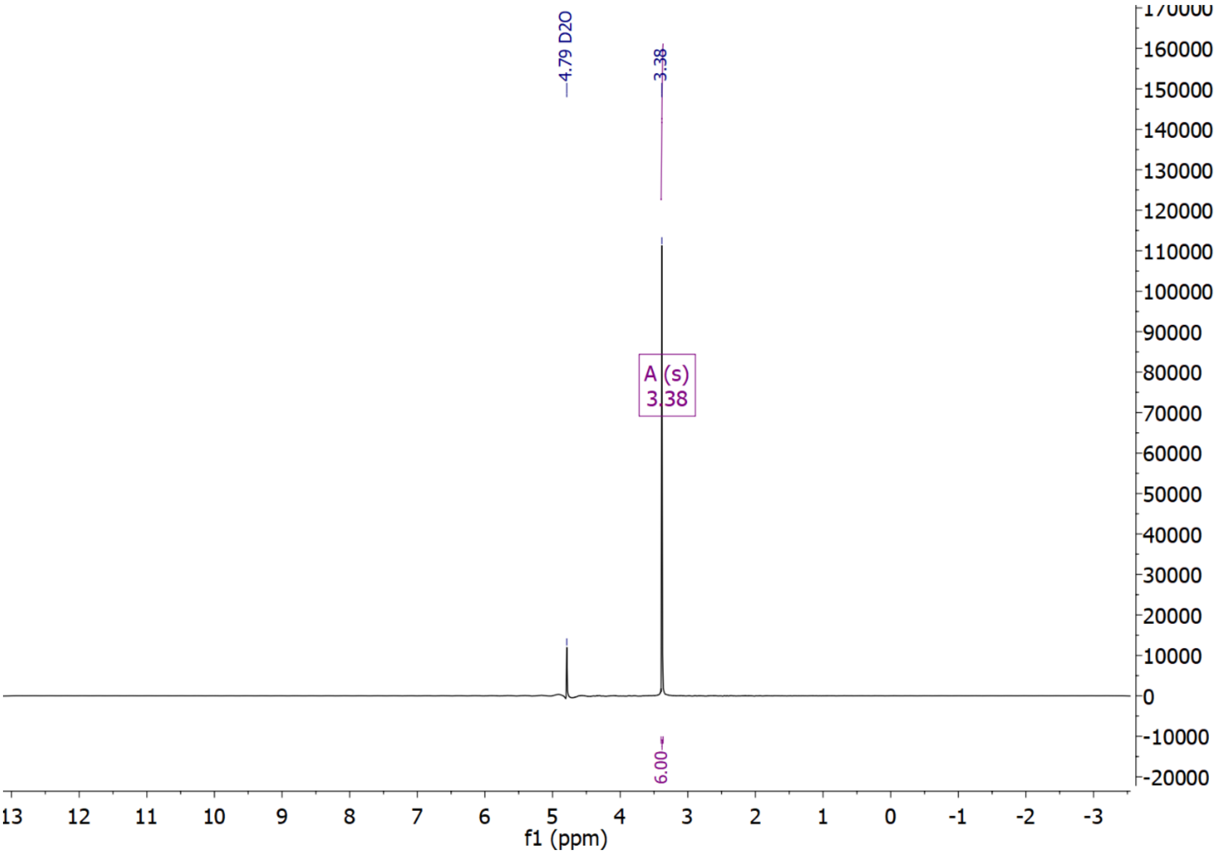


decoupled

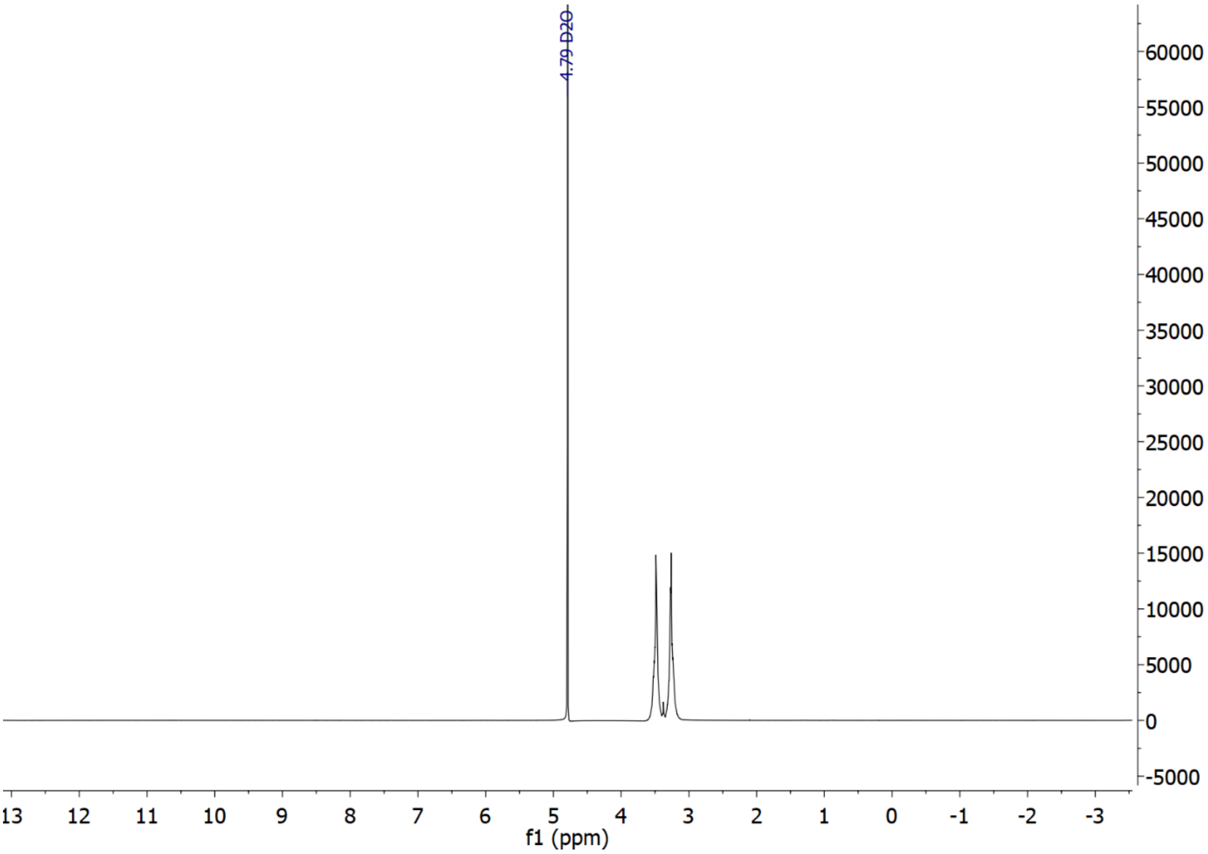

4.79 D2O
60000
55000
50000
45000
40000
35000
30000
25000
20000
15000
10000
5000
0
-5000
13 12 11 10 9 8 7 6 5 4 3 2 1 0 -1 -2 -3
f1 (ppm)


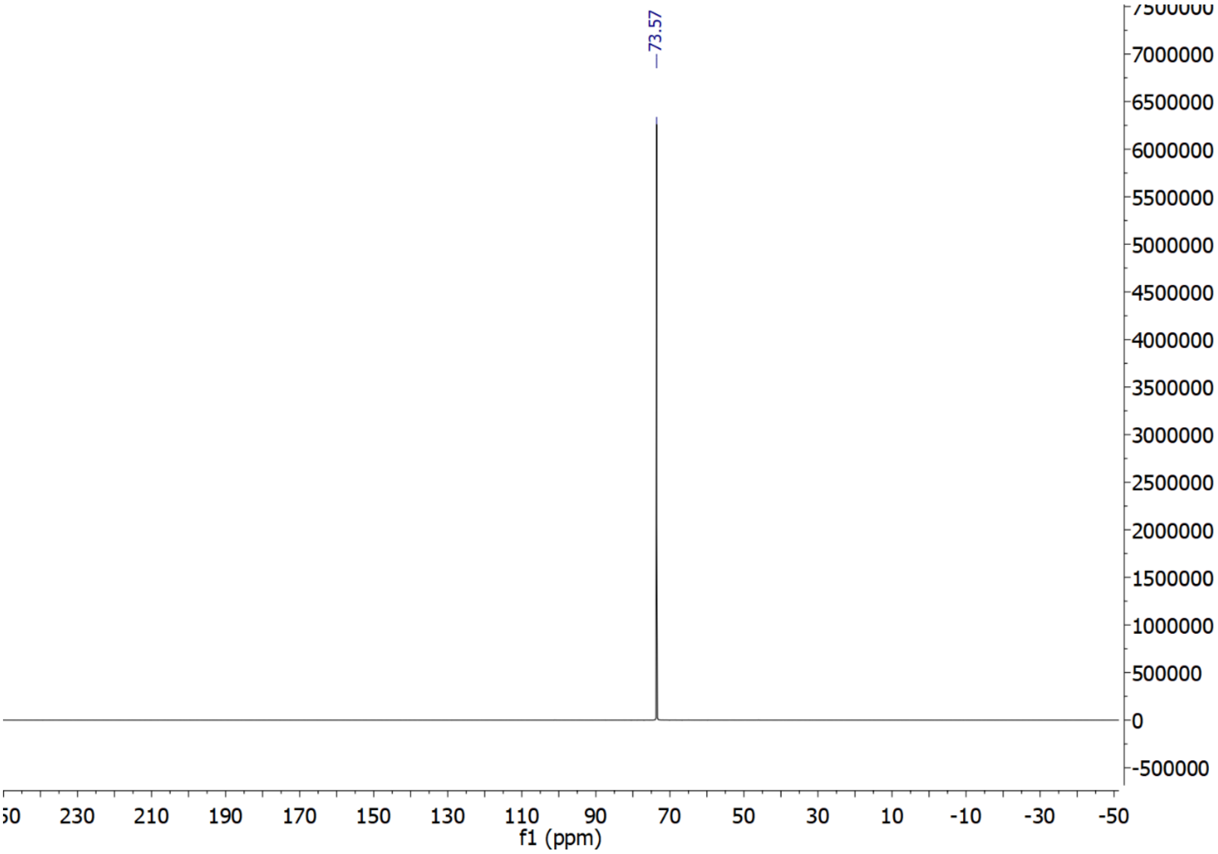

73.57
7500000
7000000
6500000
6000000
5500000
5000000
4500000
4000000
3500000
3000000
2500000
2000000
1500000
1000000
500000
0
-500000
230 210 190 170 150 130 110 90 70 50 30 10 -10 -30 -50
f1 (ppm)

**[$^{13}C_6$]*Scyllo*-inosose**

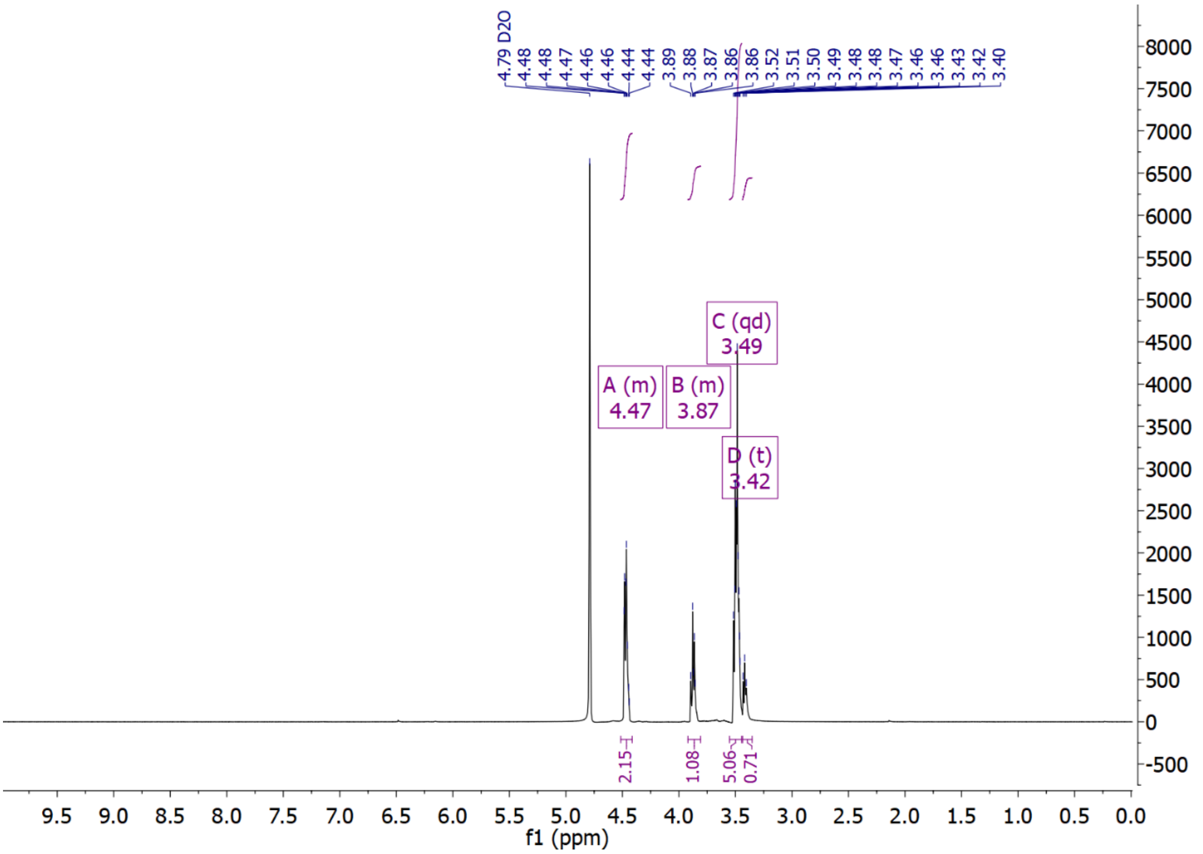


decoupled

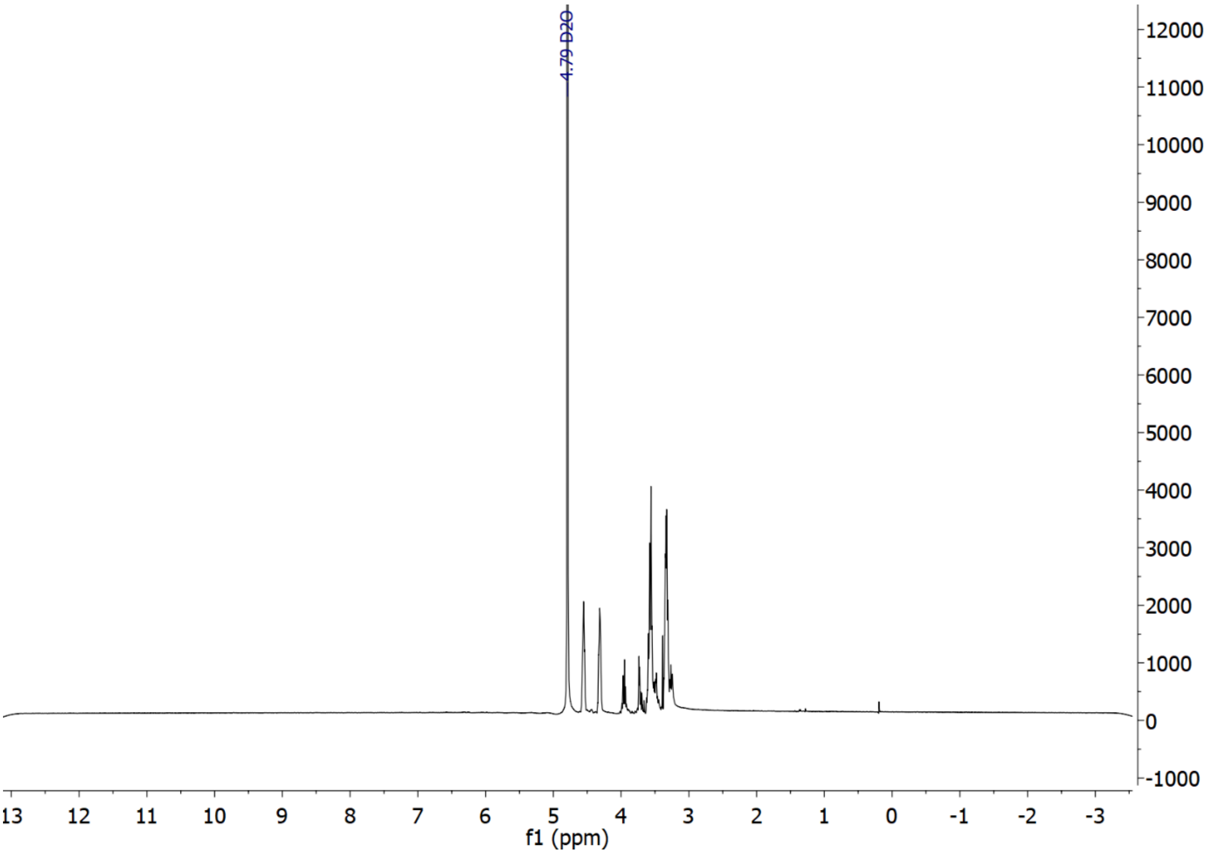

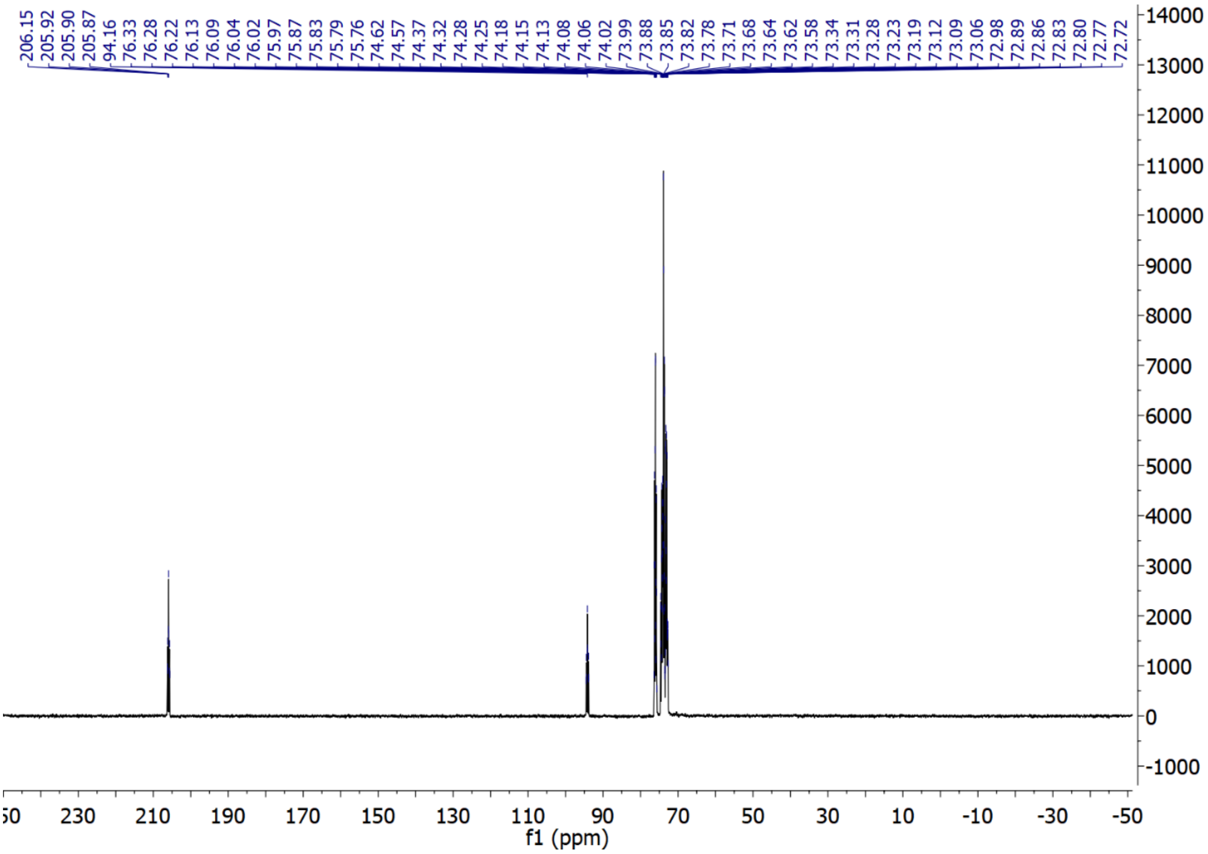
206.15
205.92
205.90
205.87
94.16
76.33
76.28
76.22
76.13
76.09
76.04
76.02
75.97
75.87
75.83
75.79
75.76
74.62
74.57
74.37
74.32
74.28
74.25
74.18
74.15
74.13
74.08
74.06
74.02
73.99
73.88
73.85
73.82
73.78
73.71
73.68
73.64
73.62
73.58
73.34
73.31
73.28
73.23
73.19
73.12
73.09
73.06
72.98
72.89
72.86
72.83
72.80
72.77
72.72
14000
13000
12000
11000
10000
9000
8000
7000
6000
5000
4000
3000
2000
1000
0
-1000
230
210
190
170
150
130
110
90
70
50
30
10
-10
-30
-50
f1 (ppm)
